# Squirrel: Reconstructing semi-directed phylogenetic level-1 networks from four-leaved networks or sequence alignments

**DOI:** 10.1101/2024.11.01.621567

**Authors:** Niels Holtgrefe, Katharina T. Huber, Leo van Iersel, Mark Jones, Samuel Martin, Vincent Moulton

## Abstract

With the increasing availability of genomic data, biologists aim to find more accurate descriptions of evolutionary histories influenced by secondary contact, where diverging lineages reconnect before diverging again. Such reticulate evolutionary events can be more accurately represented in phylogenetic networks than in phylogenetic trees. Since the root location of phylogenetic networks can not be inferred from biological data under several evolutionary models, we consider semi-directed (phylogenetic) networks: partially directed graphs without a root in which the directed edges represent reticulate evolutionary events. By specifying a known outgroup, the rooted topology can be recovered from such networks. We introduce the algorithm Squirrel (<u>S</u>emi-directed Quarnet-based Inference to <u>Re</u>construct <u>L</u>evel-1 Networks) which constructs a semi-directed level-1 network from a full set of quarnets (four-leaf semi-directed networks). Our method also includes a heuristic to construct such a quarnet set directly from sequence alignments. We demonstrate Squirrel’s performance through simulations and on real sequence data sets, the largest of which contains 29 aligned sequences close to 1.7 Mbp long. The resulting networks are obtained on a standard laptop within a few minutes. Lastly, we prove that Squirrel is combinatorially consistent: given a full set of quarnets coming from a triangle-free semi-directed level-1 network, it is guaranteed to reconstruct the original network. Squirrel is implemented in Python, has an easy-to-use graphical user-interface that takes sequence alignments or quarnets as input, and is freely available at https://github.com/nholtgrefe/squirrel.

## Introduction

Secondary contact, where diverging lineages come into contact and hybridize before continuing to diverge, is commonplace in evolution. This process is poorly described by most phylogenetic reconstruction methods which generally assume a bifurcating tree model. Secondary contact has been widely documented for diverse sets of taxa, including viruses (e.g. HIV and SARS-CoV-2, see Worobey et al. 2008; Pekar et al. 2021; Jiao et al. 2024), bacteria (e.g. Diop et al. 2022), plants (e.g. Ehrendorfer 1959; Rieseberg et al. 2003), birds (e.g. Taylor and Larson 2019), fish (e.g. Meier et al. 2019; Du et al. 2024), invertebrates (e.g. Zhang et al. 2016) and primates, including humans (e.g. Patterson et al. 2006; Green et al. 2010). Through secondary contact, introgression — the exchange of genetic material between hybridizing lineages — may occur by means of complex processes, often involving multiple rounds of backcrossing.

Evolutionary histories shaped by secondary contact can be more accurately represented by rooted phylogenetic level-1 networks than by strictly bifurcating rooted phylogenetic trees. Rooted phylogenetic level-1 networks are directed acyclic graphs that are largely tree-like in structure, describing patterns of divergence, but include localized reticulations where lineages have merged through reticulate events (see e.g. Figure 1(a) and see the Materials and Methods for a more formal definition). Application of these networks is highly desirable, but their construction is computationally intensive, and their use has remained out of reach for most biologists. Results reported here, including an efficient algorithm and software, address the challenge of building phylogenetic level-1 networks, thus offering the possibility of finding a more realistic description of biological diversity.

**Figure 1:**
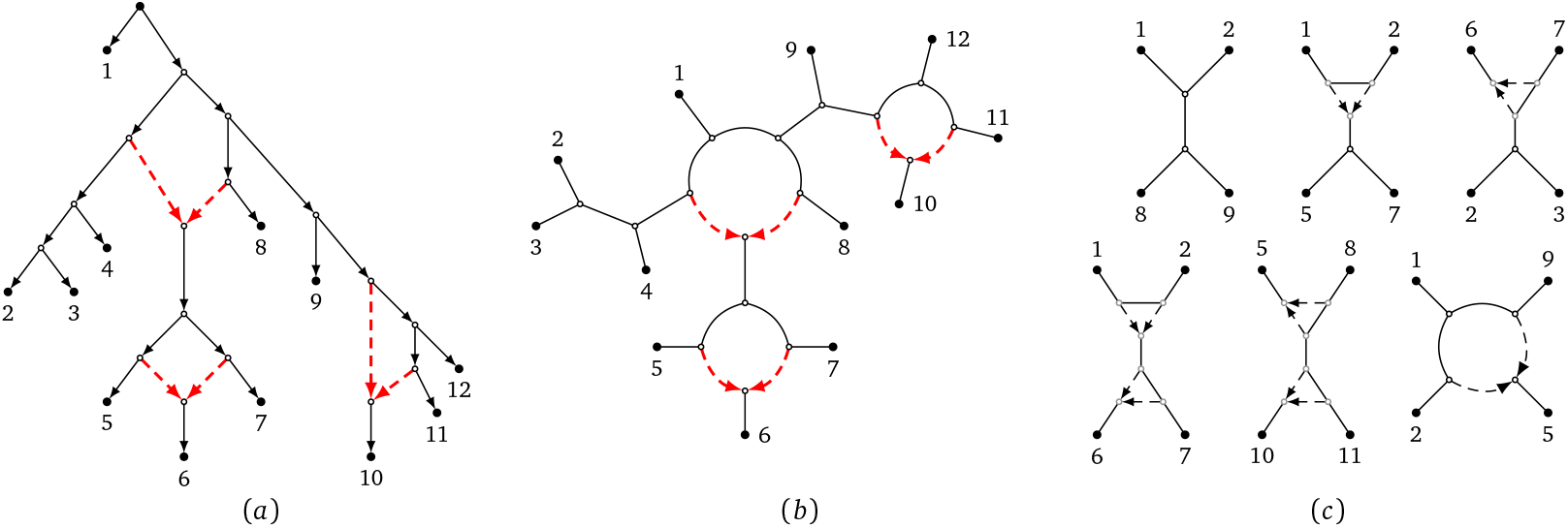
(a): A rooted phylogenetic level-1 network on 12 taxa represented by numbers 1 − 12, with the dashed reticulation edges pointing towards reticulation vertices which represent reticulate events. (b): The semi-directed topology of the rooted network, which is a triangle-free semi-directed level-1 network on 12 leaves, again with the reticulation edges dashed. This network uniquely determines the rooted network by specifying leaf 1 as an outgroup. (c): Some of the quarnets induced by the semi-directed network. When ignoring the leaf labels, these are all six possible level-1 quarnet shapes. The top left quarnet is a quartet tree, the bottom right quarnet is the only one that contains a cycle of length 4 (4-cycle), and the other four quarnets contain one or two triangles (3-cycles). The tf-quarnets (triangle-free quarnets) can be obtained from the quarnets by contracting each of the triangles to a single node. The quartet tree and quarnet with a 4-cycle are both already triangle-free.

Our results are achieved by considering *semi-directed (phylogenetic) networks* (Solís-Lemus and Ané 2016), in which there is no root and only branches representing reticulate events carry information about direction (see the Materials and Methods for a more formal definition). These networks have gained considerable interest recently (see e.g. Solís-Lemus and Ané 2016; Allman et al. 2019; Frohn et al. 2024; Kong et al. 2024; Warnow et al. 2024; Wu and Solís-Lemus 2024), as it has been shown that under certain models of evolution it is theoretically impossible to infer the root of a rooted phylogenetic network directly from data (Baños 2019; Gross et al. 2021; Xu and Ané 2023). For an example of a semi-directed level-1 network see Figure 1(b). In case an outgroup is available, this can be used to root the semi-directed network (Solís-Lemus and Ané 2016), as illustrated in Figure 1(a) and (b). Several identifiability results have been recently proven for semi-directed level-1 networks. In particular, it was shown that such networks can be theoretically recovered from data under various models of evolution (Baños 2019; Gross et al. 2021; Xu and Ané 2023). By focusing on semi-directed networks, we offer a tractable way for reconstructing phylogenetic level-1 networks.

Recently, two algebraic approaches have been introduced to construct semi-directed level-1, four-leaved networks, or *quarnets* (see Figure 1(c)): QNR-SVM (Barton et al. 2022) and an algorithm in Martin et al. (2023). These methods take as input sequence data and both employ algebraic invariants to infer quarnets under the Jukes-Cantor model (Barton et al. 2022; Martin et al. 2023) and the Kimura 2-parameter model (Martin et al. 2023). To infer evolutionary relationships for larger data sets, methods are therefore required to puzzle together such quarnets into larger networks (see e.g. Schmidt et al. (2002) and Oldman et al. (2016) for two of the earliest algorithms where this approach was used for trees and rooted networks, respectively). It is known that the quarnets coming from a semi-directed level-1 network uniquely characterize the network (Huber et al. 2024) and that theoretically they can be puzzled together efficiently to reconstruct the network (Frohn et al. 2024). However, a set of quarnets stemming from real data will unavoidably contain erroneous quarnets, thus creating the need for a more robust algorithm.

In this paper we introduce Squirrel (Semi-directed Quarnet-based Inference to Reconstruct Level-1 Networks): an efficient software tool and algorithm that builds a semi-directed level-1 network from a given full set of quarnets (that is, a *dense* set that contains one quarnet for each subset of four taxa). We complement Squir- REL with a fast heuristic method to construct quarnets from sequence data: the *δ*-heuristic (see the Materials and Methods for a formal description). Note that various existing algorithms and programs can be used to infer level-1 networks (both rooted and semi-directed) from biological data that are based on alternative approaches. For example, PhyloNet (Than et al. 2008; Yu and Nakhleh 2015), SNaQ (Solís-Lemus and Ané 2016; Solís-Lemus et al. 2017) and PhyNEST (Kong et al. 2024) are all software tools using likelihood-based algorithms operating under a coalescent model. SNaQ builds semidirected networks, whereas both PhyNEST and PhyloNet focus on rooted networks. These methods assume an upper bound on the number of reticulate events and either take gene trees (PhyloNet and SNaQ) or sequence data (PhyNEST) as input, after which they perform a potentially time-consuming search through the space of networks to optimize a likelihood criterion. On the other hand, NANUQ (Allman et al. 2019) and the recent extension NANUQ^+^ (Allman et al. 2024a) do not employ a likelihood-framework and instead use concordance factors on four-taxon subsets to produce a semi-directed level-1 network up to contracting triangles (3-cycles) and identifying the locations of reticulations in 4-cycles. This approach is faster but requires other methods to compute the input gene trees first, which itself can be a challenging step (Chifman and Kubatko 2014; Simmons and Gatesy 2015; Zhang and Mirarab 2022; Steenwyk et al. 2023). Other approaches use Bayesian methodology to construct rooted networks (e.g. SpeciesNetwork (Zhang et al. 2018a)) but are not yet able to scale to larger data sets. Lastly, Lev1athan (Huber et al. 2010) and TriLoNet (Oldman et al. 2016) take a combinatorial stance towards the network construction problem; they take as input a set of rooted three-leaf trees (Lev1athan) or rooted three-leaf networks (TriLoNet) and output a rooted level-1 network, with TriLoNet including a heuristic to generate rooted three-leaf networks from sequence data.

We now present a brief overview of how Squirrel works; a formal description of the algorithm (plus supporting figures) is given in the Materials and Methods section. As with NANUQ and to a lesser extent SNaQ, Squirrel constructs networks up to the contraction of triangles (see Figure 1(b)), thus resulting in a binary triangle-free semidirected level-1 network (i.e. a network with no cycles that contain just three vertices). Since triangles are relatively difficult to infer correctly (Gross et al. 2021), Squirrel does not use the location of any triangles in the quarnets and instead only employs *tf-quarnets* (triangle-free quarnets; see Figure 1(c)). As shown in Frohn et al. (2024), by considering tf-quarnets, we still maintain enough information to theoretically construct the complete semi-directed level-1 network up to contracting its triangles. If quarnets with triangles are given in the input, tf-quarnets are obtained by contracting the triangles. Hence, each tf-quarnet is either a quartet tree or contains a 4-cycle.

Given a dense set of weighted tf-quarnets, Squirrel first uses all of the tf-quarnets that are quartet trees to build a sequence of non-binary phylogenetic trees, using an algorithm from Berry and Gascuel (2000) and employing techniques from the QuartetJoining algorithm (Grünewald et al. 2009) that constructs phylogenetic trees from quartet trees. Within each of the non-binary phylogenetic trees in the sequence, the internal vertices with high degree are replaced by a suitable cycle. In particular, Squirrel repeatedly solves the Travelling Salesman Problem (TSP, see e.g. Bellman 1962; Held and Karp 1962) with suitably defined distances to create a cyclic ordering of the subnetworks around the cycles. This results in a sequence of candidate level-1 networks, from which Squirrel returns the one that agrees, in a well-defined sense, with most of the original tf-quarnets. If an outgroup is specified, this network can in turn be transformed into a rooted network.

We emphasize that any method that is able to create a dense set of tf-quarnets from biological data (possibly incorporating e.g. incomplete lineage sorting) could be used to generate input for Squirrel. Furthermore, Squir-REL takes into account weights the tf-quarnets might have, which can be used to model confidence or bootstrap support. Reassuringly, Squirrel is consistent in the sense that it will reconstruct the correct network if all tf-quarnets are derived from a triangle-free semi-directed level-1 network, a fact that we prove in Theorem 1 in the Materials and Methods section.

## Results

### Simulation study

Following the simulation studies for Lev1athan (Huber et al. 2010) and TriLoNet (Oldman et al. 2016), we analyze what effect noise in a set of tf-quarnets has on the performance of Squirrel. To this end, we generate 100 random triangle-free semi-directed level-1 networks for every number *n* ∈ {10, 15, 20, 25, 30, 35} of leaves (see Section B of the Supplementary Material for the generating algorithm). For each network 𝒩, the reticulation number *r*(𝒩) (i.e. the number of reticulations) is chosen uniformly at random from {0, …, ⌊ *n/*3 ⌋}. This results in a set of 600 random networks 𝒩, each inducing a set 𝒬 (𝒩) of tf-quarnets. For each network 𝒩 and each perturbation ratio *ϵ* ∈ {0, 0.01, 0.02, 0.05, 0.1, 0.2, 0.3, 0.4, 0.5}, we create a noisy set of tf-quarnets 𝒬 _*ϵ*_(𝒩) by changing the undirected underlying topology of a fraction of the tf-quarnets uniformly at random which is given by *ϵ*. Then, if this creates a 4-cycle, we pick a random location for the reticulation. We use this scheme for the creation of noise to prevent 4-cycles from only changing their reticulation and keeping their circular ordering. Such a perturbation will barely influence the output of the algorithm, since reticulations of 4-cycle tf-quarnets are only used to determine the location of reticulations in 4-cycles of the final networks. The resulting 5400 = 600 · 9 sets of unweighted tf-quarnets 𝒬 _*ϵ*_(𝒩) are used as input for Squirrel. The average computation times ranged from below a second for the networks with the fewest leaves to below two minutes for the networks with 35 leaves.

To measure how well Squirrel reconstructs the original networks from these noisy tf-quarnet sets we compute two similarity scores for every input network and 𝒩 output network ℳ. The first score is the *tf-quarnet consistency score* (modeled after a similar score in Huber et al. (2010) and Oldman et al. (2016)) which is defined as

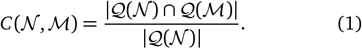

This score measures what fraction of the tf-quarnets induced by 𝒩 are also induced by the constructed network ℳ. We also consider its symmetric counterpart: the *tf-quarnet symmetric consistency score*, defined as

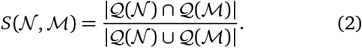

Both scores are always in the interval [0, 1] and attain a value of 1 if and only if 𝒩 = ℳ, which follows from Frohn et al. (2024). The boxplots in Figure 2 show the distribution of the two scores for different perturbation ratios *ϵ* and leaf set sizes *n*. As expected, both scores decrease for larger values of *ϵ*. However, the decrease seems fairly limited, with both consistency scores averaging above 0.91 even for sets containing only 50% of the original tf-quarnets.

**Figure 2:**
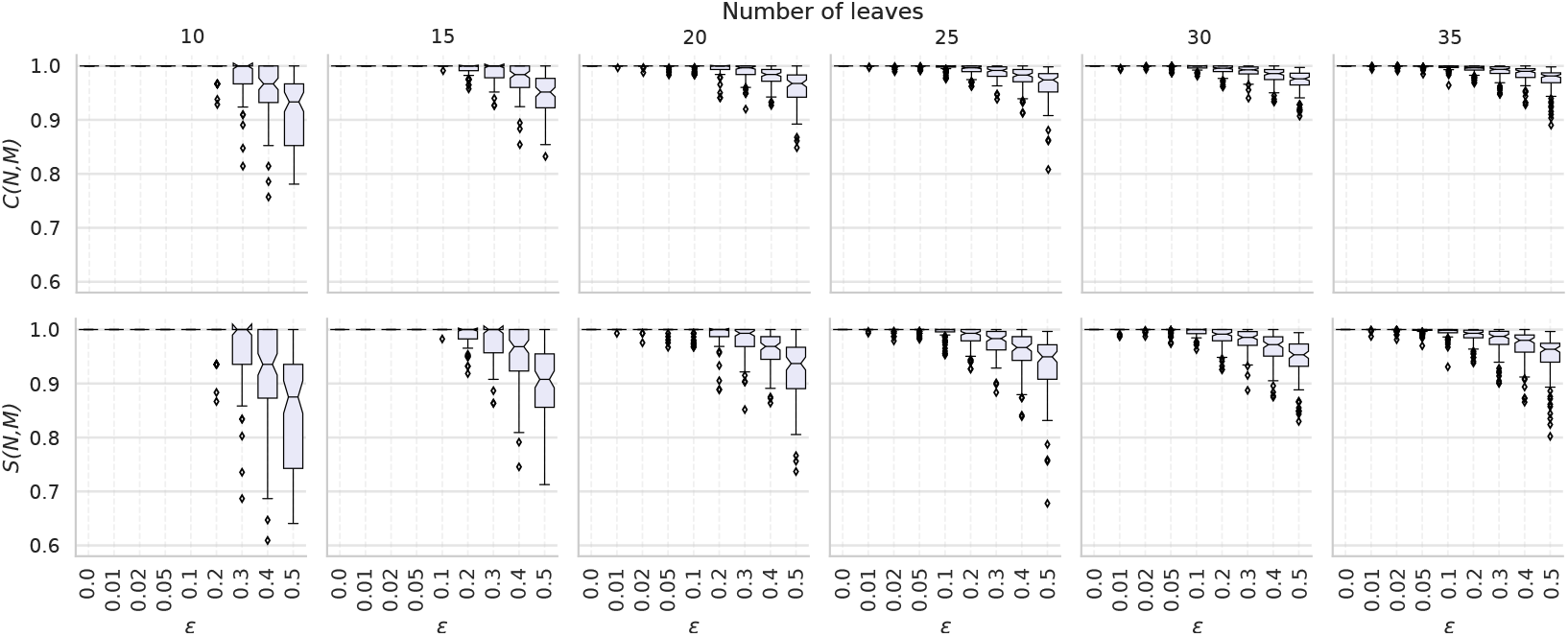
Boxplots showing the spread of *C*- and *S*-scores between the input network 𝒩 and output network 𝒩 ℳ, when applying Squirrel to sets of tf-quarnets with leaf set sizes *n* and perturbation ratios *ϵ*. The boxplots show the quartiles of the data and its outliers. A single outlier in the case of *n* = 10 and *ϵ* = 0.5 has a *C*- and *S*-score below 0.6 and is omitted from the figure for clarity.

To investigate in what way noise in a set of tf-quarnets influences the structure of the reconstructed networks, we compute the difference in the reticulation numbers *r*(𝒩) − *r*(ℳ) between the input networks 𝒩 and output networks ℳ. The boxplots in Figure 3 show the result of this experiment, again for different values of *ϵ* and *n*. Up to a value of *ϵ* = 0.1, Squirrel reconstructs networks with the correct reticulation number in almost all cases. For higher values, the differences are more spread out, while the average difference slowly becomes positive. Thus, it seems that Squirrel slightly favors networks with fewer reticulations for high values of *ϵ*, although the average absolute differences remain below a reasonably small 1.5. A possible explanation could be that by not considering triangles in the quarnets, the signal in the data indicating reticulate events is weakened.

**Figure 3:**
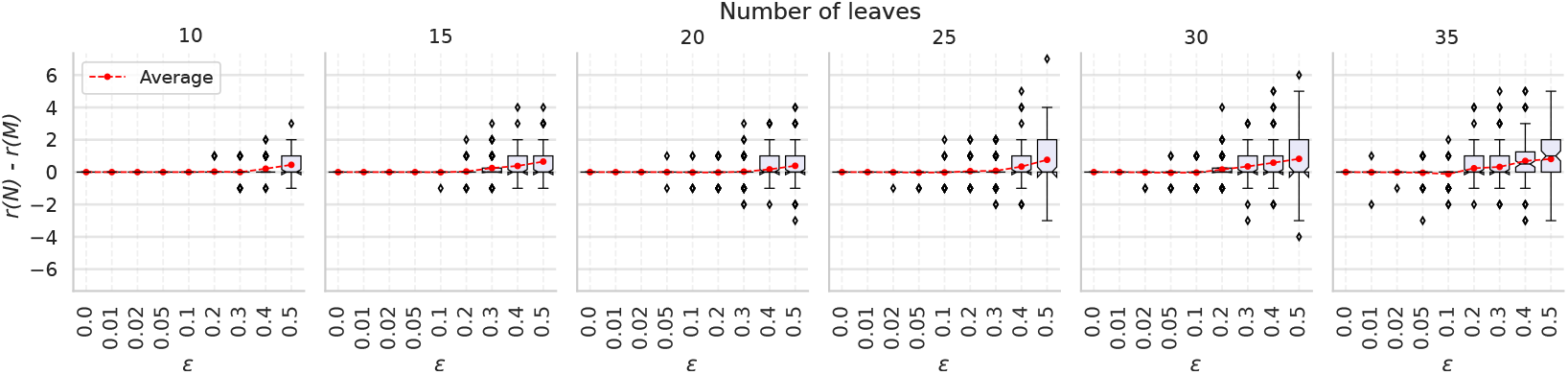
Boxplots showing the variation of the difference in reticulation number *r*(𝒩) − *r*(ℳ) of the input network 𝒩 and output network ℳ, when applying Squirrel to sets of tf-quarnets with leaf set sizes *n* and perturbation ratios *ϵ*. The boxplots show the quartiles of the data, its outliers and the averages in red.

We also perform a study with simulated nucleotide sequences to test the performance of the *δ*-heuristic combined with Squirrel, using a similar approach to the simulations presented in Holland et al. (2002) and Oldman et al. (2016). For each of our 600 previously generated networks, we simulate one multiple sequence alignment (MSA) for every sequence length *k* ∈ {1 kbp, 10 kbp, 100 kbp, 1 Mbp} as follows. Briefly, we first root every semi-directed network 𝒩 uniformly at random on some edge (making sure that it is a valid root-location) to create a rooted phylogenetic network. We then use the software tool Seq-Gen (Rambaut and Grass 1997) to simulate MSAs of equal length along all displayed trees of the rooted phylogenetic network under the K2P model with transition-transversion bias 4 (as in Holland et al. 2002; Oldman et al. 2016). The MSAs of the displayed trees are then concatenated to create one MSA with the desired length *k*. Since our *δ*-heuristic treats every site of the MSA independently, this way of generating MSAs is asymptotically equivalent to generating MSAs under the K2P network-based Markov model with reticulation parameters of 0.5 (see e.g. Gross et al. 2021).

The branch lengths (i.e. the expected number of substitutions along each edge) that are used for the simulations are determined as follows. Given an edge (*u, v*) of one of the rooted phylogenetic networks, we let *p*_(*u,v*)_ be the average length (in terms of number of edges) of all unique paths from the root to any leaf that contain the edge (*u, v*). Then, we assign the edge (*u, v*) a branch length of 0.3*/p*_(*u,v*)_, which ensures that every path in the network from a root to a leaf roughly has a total length of 0.3, as is the case in the simulations by Holland et al. (2002) and Oldman et al. (2016).

We then use the 2400 = 600 · 4 simulated MSAs as input for our *δ*-heuristic to construct dense sets of weighted tf-quarnets, which are in turn used to construct semi-directed networks with Squirrel. As before, we compare every constructed semi-directed network *ℳ* with the original semi-directed network 𝒩 in terms of *C*-score, *S*-score and difference in reticulation number *r*(𝒩) − *r*(ℳ). The results are depicted in Figure 4 and Figure 5, respectively. We observe that both consistency scores increase as the sequence length changes from 1 kbp to 10 kbp. Additionally, both the average and the variation of the difference in reticulation number decrease. Interestingly, the increase of the sequence length from 10 kbp to 100 kbp or 1 Mbp does not seem to have much further effect. As was the case in our previous experiment, an increase in the number of leaves *n* of the original semi-directed network improves the two considered consistency scores, yet also results in a greater spread of the difference in reticulation number between the original and constructed network. The latter point can be explained by the fact that smaller networks simply allow for fewer reticulations, thus also bounding the largest possible difference in reticulation number.

**Figure 4:**
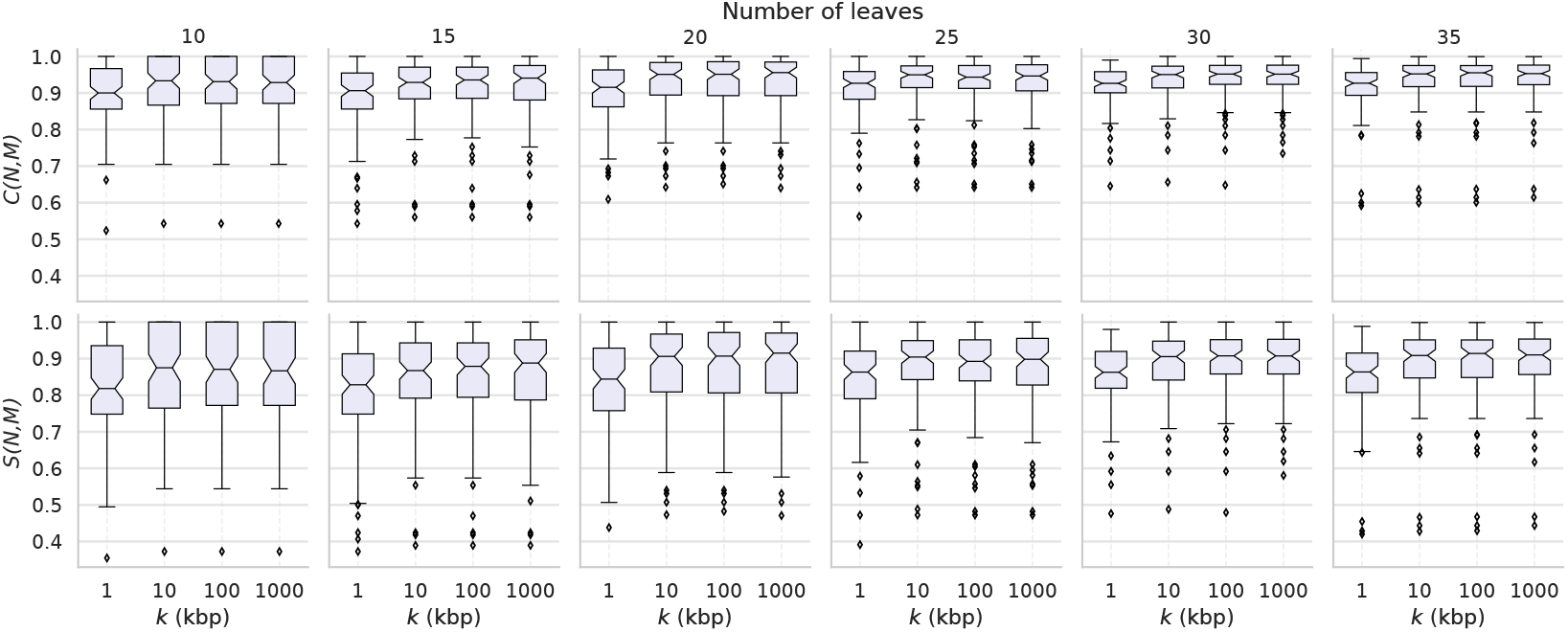
Boxplots showing the spread of *C*- and *S*-scores between the input network 𝒩 and output network ℳ, when applying the *δ*-heuristic and Squirrel to MSAs with leaf set sizes *n* and sequence lengths *k*. The boxplots show the quartiles of the data and its outliers.

**Figure 5:**
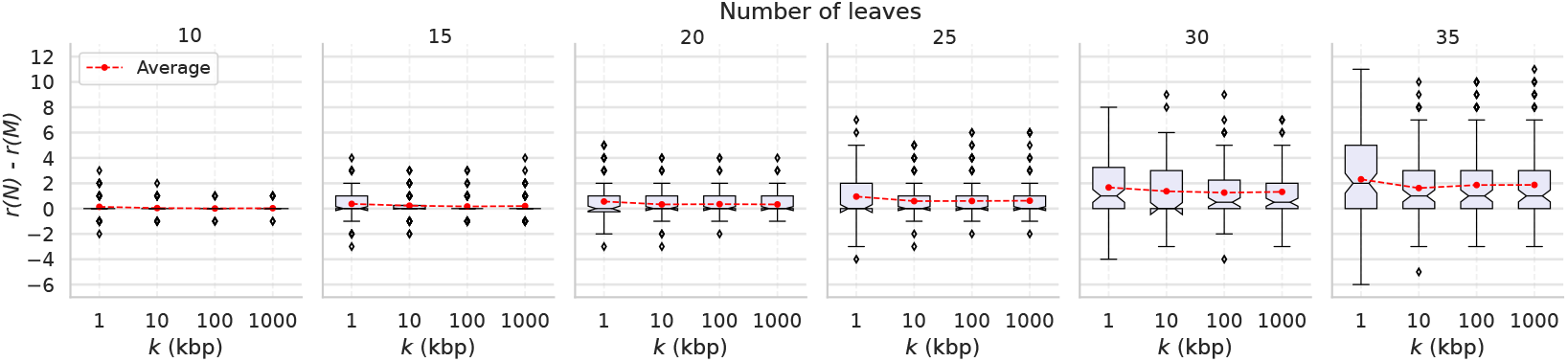
Boxplots showing the variation of the difference in reticulation number *r*(𝒩) − *r*(ℳ) of the input network 𝒩 and output network ℳ, when applying the *δ*-heuristic and Squirrel to MSAs with leaf set sizes *n* and sequence lengths *k*. The boxplots show the quartiles of the data and its outliers.

### Biological data

To illustrate the applicability of Squirrel to biological data, we consider three data sets on groups of taxa with evidence of secondary contact in their evolutionary histories: a large set of tf-quarnets generated with the MML algorithm from Martin et al. (2023) (named after the authors), a short multiple sequence alignment on few taxa from Salemi and Vandamme (2003), and a long multiple sequence alignment on many taxa from Vanderpool et al. (2020).

### Xiphophorus

We first test the applicability of Squirrel to a set of tf-quarnets that was generated with the MML algorithm (Martin et al. 2023). For each four-taxon subset, this algorithm creates a ranking of the possible 4-cycles according to some scoring criterion (with the lowest score being the best). Based on the scores it either detects a quartet tree (which we give a weight of 1), or it chooses the best 4-cycle, which we give a weight of min(1, *s*_2_*/s*_1_ − 1), where *s*_1_, *s*_2_ are the two lowest (and thus best) scores. In this manner we take into account how close the scores for the two best scoring 4-cycles are.

The data set we consider contains 14,950 weighted tf-quarnets on a set of 25 swordtail fish and platyfish (genus *Xiphophorus*) and the single outgroup *Pseudoxiphophorus jonesii*. This genus has been widely studied and much evidence has been presented for widespread hybridization within the genus (see e.g. Rosenthal et al. 2003; Culumber et al. 2011; Cui et al. 2013; Kang et al. 2013; Schumer et al. 2013; Solís-Lemus and Ané 2016, and the references therein), making it difficult to capture the full evolutionary history. Traditionally, the genus is divided into four major lineages: northern swordtails, southern swordtails, northern platyfishes and southern platyfishes (Meyer et al. 2006; Cui et al. 2013). The best network generated by Squirrel (taking less than two minutes) had a weighted tf-quarnet consistency score of 0.974 and is shown in Figure 6. However, many of the other candidate networks had scores that were very close to the score of the best scoring network.

**Figure 6:**
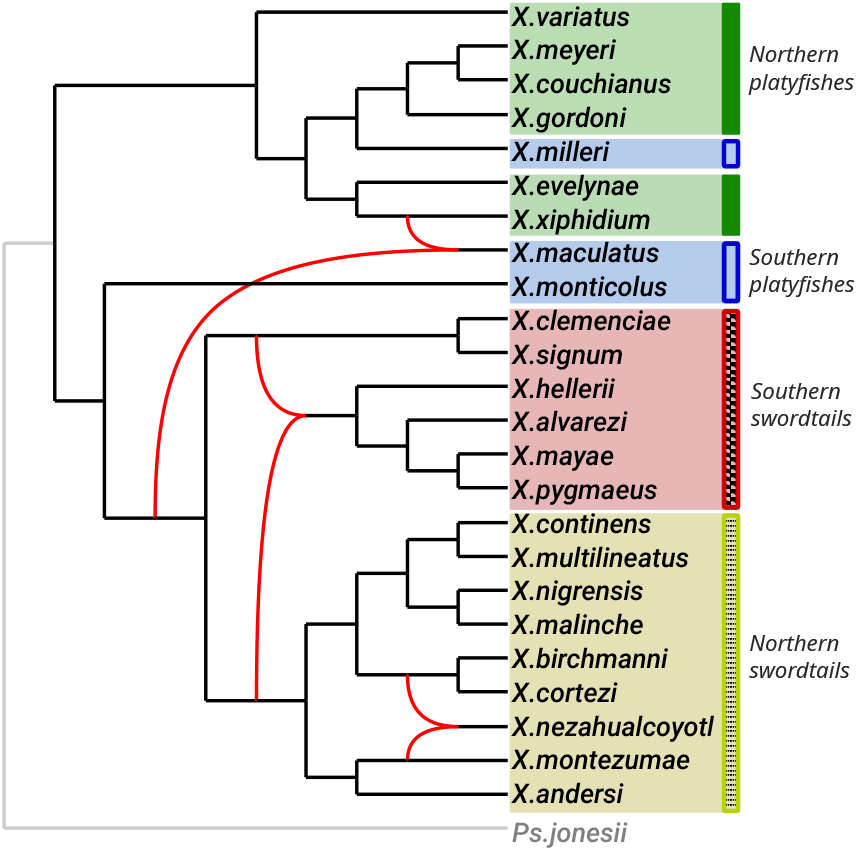
Phylogenetic network inferred by Squirrel from a dense set of weighted tf-quarnets on the genus *Xiphophorus* (generated from a multiple sequence alignment with the MML algorithm from Martin et al. (2023)). The four major lineages are indicated by the different shaded areas. The reticulation edges are curved, while the edges leading to the outgroup *Pseudoxiphophorus jonesii* are in grey.

Since the weighted tf-quarnet consistency score measures how consistent the network is with the tf-quarnets, taking their weights into account (see eq. (3) in the Materials and Methods), it should be noted that a weighted consistency score close to 1 does not necessarily imply a close to 100% level of confidence that the network is correct. Instead, it reflects whether the quarnets with high weight (i.e. high confidence in their correctness) are consistent with the constructed network, making it most useful as a relative measure to assess if there is a clear best network or if multiple networks perform similarly well. In contrast, the unweighted consistency score (see eq. (1)) can be more easily interpreted as an absolute measure of performance, but it may discard useful information about quarnet confidence if such information is available. A more statistically sound way to generate weights for the tf-quarnets inferred with the MML algorithm from Martin et al. (2023) (similar to the bootstrap support in Barton et al. (2022)) would possibly increase the confidence of Squirrel in a single best network. Hence, we would welcome further research efforts into computing confidence scores for inferred tf-quarnets which can be used as input weights for Squirrel.

The constructed network clearly divides the three major *Xiphophorus* clades (northern swordtails, southern swordtails and platyfishes) but similar to other studies (Meyer et al. 2006; Cui et al. 2013) intertwines northern and southern platyfishes. Our network has one reticulation edge involving an ancestor of both the northern and the southern swordtails. Another reticulate event places the northern swordtail *X*.*cortezi* both as a sibling of *X*.*nezahualcoyotl* and of the clade (*X*.*malinche, X*.*birchmanni*). This reticulate event aligns with previous work in Cui et al. (2013), where the precise placement of *X*.*cortezi* within this subset of the species (including *X*.*montezumae*) was also uncertain and depended on the inference methods used. Furthermore, one of the subtrees displayed in our network for this subset of the species (i.e. the subtree that includes *X*.*montezumae*) is the same as the subtree of the network inferred by SNaQ (Solís-Lemus and Ané 2016; Solís-Lemus et al. 2017). The last reticulate event involves the southern platyfish *X*.*maculatus*, for which Cui et al. (2013) report difficulties placing it in the mitochondrial DNA tree. Judging from the many different inferred networks and possible reticulate events (see again Rosenthal et al. 2003; Culumber et al. 2011; Cui et al. 2013; Kang et al. 2013; Schumer et al. 2013; Solís-Lemus and Ané 2016), capturing the evolutionary history of the complete genus as a level-1 network might be too much to ask for because the truth may not be level-1. As an example, evolutionary histories containing many hybridization events between more distantly related species (such as horizontal gene transfer) can not always be captured well by a level-1 network, since such events often result in complex networks with many nested reticulation events (see e.g. Soucy et al. 2015, Fig. 5).

### HIV

We now consider a multiple sequence alignment (MSA) of the HIV-1 virus data set containing 9 sequences of length 9,953 bp which first appeared in Salemi and Vandamme (2003). This data set is well-studied (Lemey et al. 2009; Huber et al. 2010; Oldman et al. 2016) and contains sequences of the HIV-1 M-group subtypes *A, B, C, D, F, G, H* and *J* as well as a sequence for *KAL153* which is believed to be a recombinant of subtypes *A* and *B* (see Lemey et al. 2009, Ch. 16). We use our *δ*-heuristic (formally described in the Materials and Methods) to obtain a weighted set of tf-quarnets from the MSA and then apply Squirrel to construct a network, which we root using the outgroup *C* (as in Salemi and Vandamme 2003; Huber et al. 2010). The *δ*-heuristic and Squirrel constructed a clear best scoring network (shown in Figure 7(a)) with a weighted tf-quarnet consistency of 0.58 within one second.

**Figure 7:**
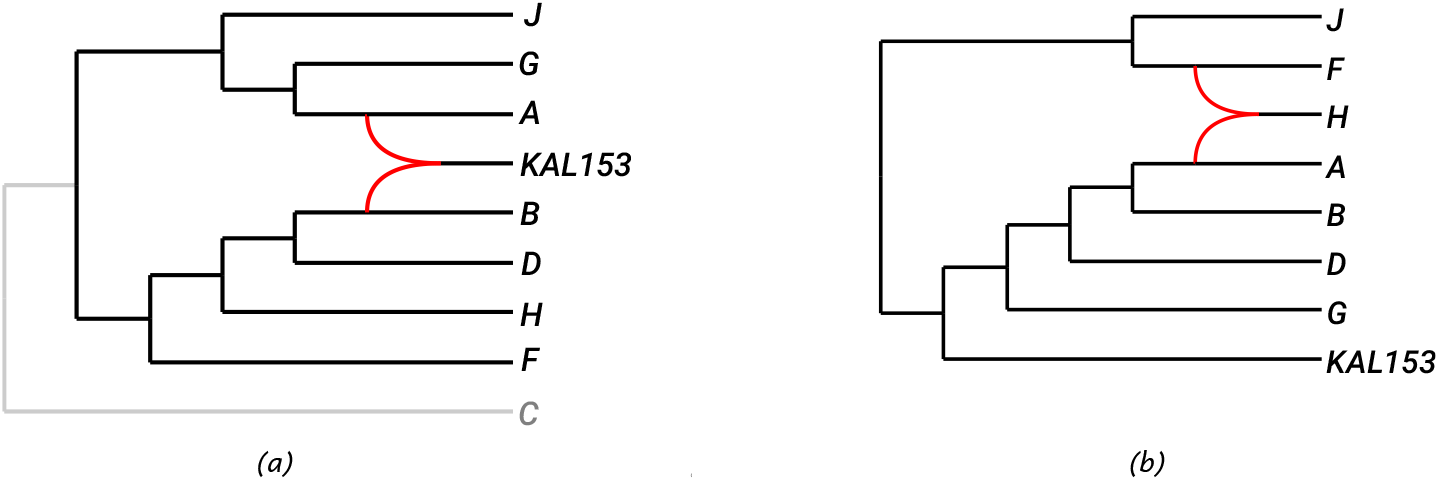
(a): Phylogenetic network inferred by Squirrel (using the *δ*-heuristic to create tf-quarnets) from a multiple sequence alignment of the HIV-1 data set under consideration. The reticulation edges are curved, while the edges leading to the outgroup *C* are in grey. (b): Phylogenetic network inferred by TriLoNet (Oldman et al. 2016) on the same HIV-1 data set (without the outgroup *C*), again with curved reticulation edges.

Indeed, Squirrel, combined with the *δ*-heuristic, is able to identify *KAL153* as a recombinant of subtypes *A* and *B*, agreeing with the analysis in (Lemey et al. 2009, Ch. 16). This compares favourably to TriLoNet (Oldman et al. 2016), where the subtype *H* was identified as a recombinant (see the constructed network in Figure 7(b)). Lev1athan (Huber et al. 2010) was able to identify *KAL153* as a recombinant, but it relies on other algorithms to make the step from sequences to gene trees.

### Primates

To investigate the performance of Squirrel and the *δ*-heuristic on data sets with many taxa and long sequences, we consider an MSA from Vanderpool et al. (2020) of length 1,761,114 bp that contains concatenated sequences for 26 primate species, 2 closely related nonprimate species and the outgroup *Mus musculus*. We first apply the *δ*-heuristic to the MSA to obtain a set of 23,751 weighted tf-quarnets. Subsequently, we use Squirrel (specifying *Mus musculus* as the outgroup to root it) and obtain the tree in Figure 8(a) after a few minutes on a standard laptop. The tree coincides exactly with the species tree obtained in Vanderpool et al. (2020) using the gene tree based algorithm ASTRAL III (Zhang et al. 2018b), while largely agreeing with two previously inferred phylogenies (Perelman et al. 2011; Springer et al. 2012). The weighted tf-quarnet consistency score of the tree is 0.995, but some of the other generated candidate networks (which contain reticulations) have scores within 0.003 from this best value, suggesting that reticulate events might have occurred.

**Figure 8:**
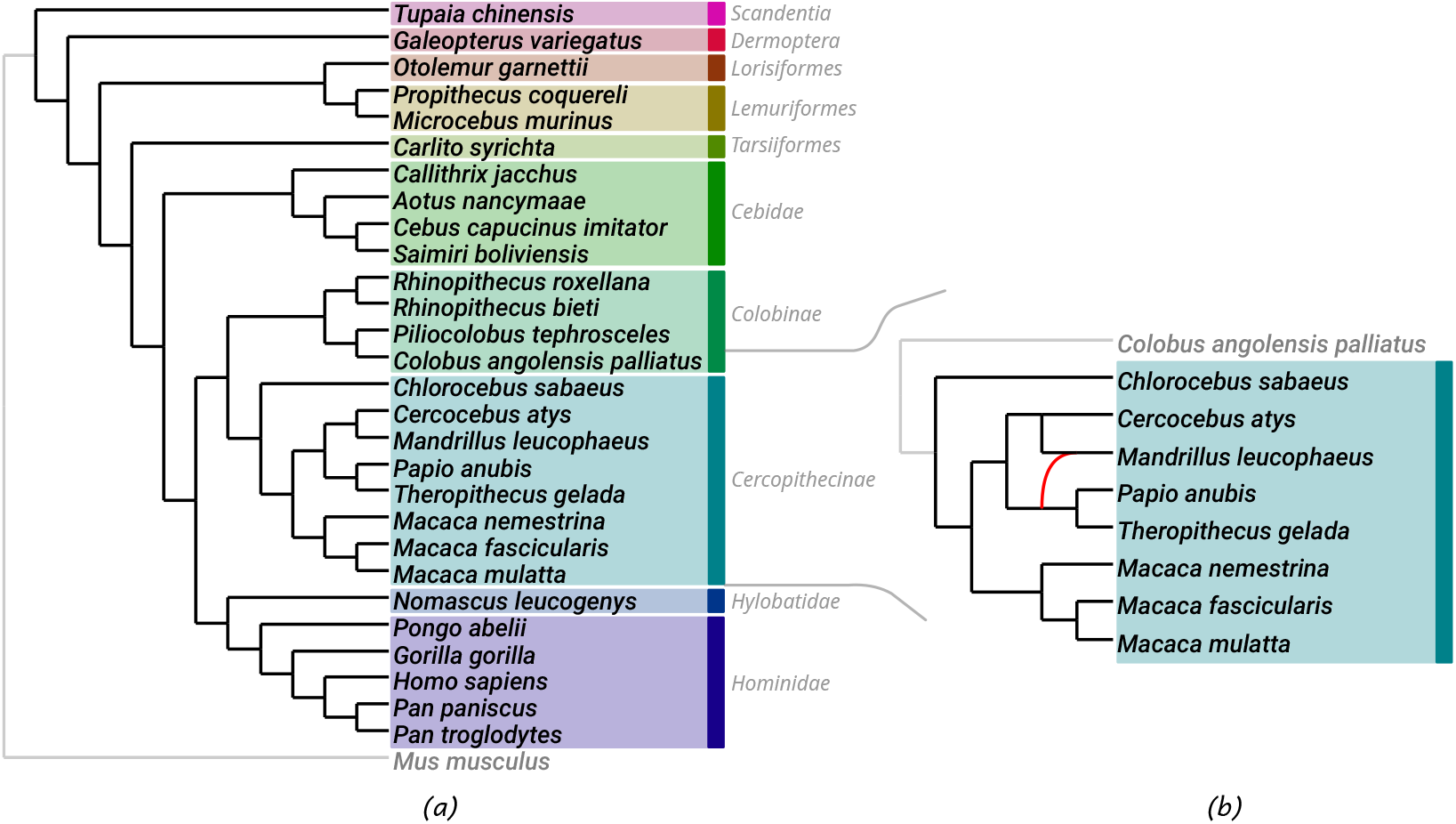
(a): Phylogenetic tree inferred by Squirrel (using the *δ*-heuristic to create tf-quarnets) from a multiple sequence alignment of the primate data set under consideration, with the edges leading to the outgroup *Mus musculus* in grey. The different shaded areas indicate different taxonomical groups as they appear in Vanderpool et al. (2020). The two non-primate species are *Tupaia Chinensis* and *Galeopterus variegatus*. (b): In conjunction with the *δ*-heuristic to create tf-quarnets, Squirrel inferred two networks with very close weighted tf-quarnet consistency scores from the considered multiple sequence alignment of the subfamily of *Cercopithecinae* (using *Colobus angolensis palliatus* as outgroup). One of them is the depicted network and the other is the phylogenetic tree obtained from that network by ignoring the curved reticulation edge.

We investigate this further by looking only at the 8 primates in the *Cercopithecinae* subfamily, for which Vanderpool et al. (2020) have demonstrated possible reticulate events. Combining the *δ*-heuristic and Squirrel we generated a set of candidate networks for these 8 species and the outgroup *Colobus angolensis pallatus*. Two of the networks had a much higher score than the others and they only differed from each other by the addition of a reticulation edge. In particular, the second best scoring network (shown in Figure 8(b)) had a score of 0.956, while the best scoring network was the subtree of the original network with score 0.974 (also shown in Figure 8(b), by ignoring the curved reticulation edge). The *blobtree* of the network (obtained by contracting the cycle into a single node) exactly matches one of the blobtrees inferred with TINNiK (Allman et al. 2024b). The particular reticulate event we found was not reported in Vanderpool et al. (2020). However, our reticulate event might be more probable since it is between species in the same continent (Africa), while the study by Vanderpool et al. (2020) mentions possible reticulate events between species on different continents (Asia and Africa). Lastly, Vanderpool et al. (2020) found evidence for a “complex pattern of ancient introgression” (p. 14) within the subfamily and state that roughly 40% of the species within the subfamily are known to hybridize (Tung and Barreiro 2017), which suggests that the true nature of the subfamily might not be well-represented by a level-1 network. This is further supported by the fact that the analysis done in Vanderpool et al. (2020) with PhyloNet (Than et al. 2008; Yu and Nakhleh 2015) and SNaQ (Solís-Lemus and Ané 2016; Solís-Lemus et al. 2017) also gave ambiguous results, while PhyNEST (Kong et al. 2024) yet again concludes with a different network.

The *Cercopithecinae* subfamily (again with outgroup *Colobus angolensis pallatus*) also featured in Barton et al. (2022) in the context of using the QNR-SVM algorithm for inferring quarnets from a data set. The reason for restricting to a subset was stated as the lack of an algorithm that puzzles together many quarnets. Instead, the authors puzzle them together by hand to obtain a network with a single reticulation that induces 81% of the well-supported quarnets. Using their quarnet weighting scheme, Squirrel was able to identify a tree inducing 85% of the well-supported quarnets. (Here, we used a variation of Squirrel that takes into account the triangles of the quarnets to choose the best scoring network, instead of the default of just focusing on the tf-quarnets). Therefore, Squirrel might be a viable tool to puzzle together quarnets obtained with an algorithm such as QNR-SVM, while still being able to scale to larger data sets unfit for resolving conflicting quarnets by hand.

## Discussion

We have introduced Squirrel: a combinatorially consistent algorithm that can puzzle together a dense set of quarnets to create a semi-directed level-1 network. In addition, when combined with the model-based method QNR-SVM (Barton et al. 2022) or the MML algorithm (Martin et al. 2023) for inferring quarnets, Squirrel provides a method to create a level-1 network directly from sequence data. To the best of our knowledge, Squirrel is one of the first methods that allows the construction of semi-directed level-1 networks from biological data using collections of quarnets. The only other approaches we are aware of that use quarnet information are NANUQ (Allman et al. 2019) and the recently presented NANUQ^+^ (Allman et al. 2024a). Although NANUQ^+^ uses a similar distance-based strategy to Squirrel to expand the cycles in a network, both NANUQ and NANUQ^+^ take as input a collection of gene trees, rather than a dense set of quarnets or a sequence alignment.

Any method that creates a dense set of quarnets from biological data could be used as input for Squirrel. In particular, if such a method is statistically consistent under some model (possibly incorporating e.g. incomplete lineage sorting), the combinatorial consistency of Squirrel ensures that the combined inference is consistent as well. Furthermore, Squirrel could in principle be combined with methods that may not scale well to larger taxa sets but are still able to construct partial semi-directed level-1 networks (containing some but not all of the studied taxa) from biological data. Indeed, as with supertree methods, partial networks on larger sets of taxa could be converted to quarnets for Squirrel by restricting those partial networks to four taxa. This would require a rule to decide what to do in case partial networks overlap on more than four taxa and they induce conflicting quarnets. Hence, a possible direction for future research would be adapting Squirrel to work with non-dense sets of quarnets which could contain any number of quarnets for each subset of four taxa.

Using the *δ*-heuristic, Squirrel is able to quickly construct a level-1 network directly from sequence data. Our sequence simulations show that the *δ*-heuristic is likely not statistically consistent under the tested K2P model. In particular, an increase in sequence length beyond 10 kbp does not give a visible improvement under our simulation settings, which one would expect for a statistically consistent quarnet inference method. Despite the lack of a statistical basis of the *δ*-heuristic, it already shows promising similarity scores for multiple sequence alignments with a length of 1 kbp when combined with Squirrel. Furthermore, a major advantage is its speed. As an example, this approach was able to construct a network with 29 taxa from a multiple sequence alignment of length 1.7 Mbp within a few minutes on a standard laptop (see the Results section). Hence, we do not see the *δ*-heuristic (combined with Squirrel) as an alternative for known model-based methods, but rather as a complementary tool. For one, this approach can be used to generate reasonable starting networks for the time-intensive search through the network-space of likelihood-based methods (such as Phy-loNet (Than et al. 2008; Yu and Nakhleh 2015), SNaQ (Solís-Lemus and Ané 2016; Solís-Lemus et al. 2017) and PhyNEST (Kong et al. 2024)). On the other hand, it can be used to quickly gain insight into sequence data without the need to first infer gene trees with a different tool, as is the case for NANUQ (Allman et al. 2019), which requires many accurate gene trees to make a good estimate of the concordance factors.

With the increasing availability of genome and transcriptome data, biologists are also likely to explore the reconstruction of separate phylogenetic networks for multiple sets of short orthologous sequences. Rapid construction of such networks for the same set of taxa across different sets of orthologues opens up the possibility for comparative analyses. A possible research direction in this area would be to combine Squirrel’s speed for constructing semidirected level-1 networks with the tf-quarnet consistency score or the recently introduced dissimilarity measure for semi-directed networks that generalizes the widely-used Robinson-Foulds distance for phylogenetic trees (Maxfield et al. 2024), which would permit the rapid comparison of networks computed for different sets of orthologues. It also leads to the interesting problem of finding a consensus of a collection of semi-directed networks, which to our best knowledge has not yet been addressed in the literature. One approach to this problem could be to treat it as a supernetwork question where all input networks have the same leaf set, and use the approach suggested earlier in this section.

Our simulations indicate that Squirrel can construct networks closely resembling an underlying network in terms of tf-quarnets, even if many of the tf-quarnets are wrongly inferred. In particular, both of the considered consistency scores average above 0.91 even for sets containing only 50% of the original tf-quarnets. This is a significant improvement compared to a similar experiment to the triplet*/*trinet-based Lev1athan and TriLoNet algorithms, where the *trinet consistency score* (the rooted three-leaf network analogue of our tf-quarnet consistency score) dropped below 0.5 for sets still containing 75% of the trinets (Oldman et al. 2016). These results can be considered as evidence that Squirrel is able to construct networks with a high topological resemblance to the original network in terms of tf-quarnets, even for a high percentage of incorrect tf-quarnets. As mentioned in the Results section, even though the tf-quarnets are theoretically enough to construct a triangle-free semi-directed level-1 network, in practice, contracting the triangles might somewhat weaken the signal of reticulation events. Note that theoretically (that is, when all quarnets come from a single network with *n* leaves) only 𝒪 (*n* log *n*) tf-quarnets are required to reconstruct the network, instead of the full set of 𝒪 (*n*^4^) tf-quarnets (Frohn et al. 2024). Thus, even sets with many incorrect tf-quarnets might still hold enough information to reconstruct the original network. This could also explain why a higher number of leaves seems to have a positive effect on the similarity score: 𝒪(*n* log *n*) grows slower than 𝒪 (*n*^4^), so the fraction of tf-quarnets necessary to reconstruct a network decreases when *n* grows.

Although several methods can construct semi-directed level-1 networks, the assumption that a network is level-1 might be too restrictive in many cases for biological data. A major breakthrough would be to develop a practical algorithm that is able to construct networks that are more complex than level-1 networks. Some theoretical results have already appeared towards tackling this problem. For example, it is known that semi-directed level-2 networks are uniquely encoded by the quarnets they induce (Huber et al. 2024). In addition, under several models, the circular ordering around the blobs of outerlabeled planar networks (a class of semi-directed networks more general than semi-directed level-1 networks) is also shown to be identifiable (Rhodes et al. 2025). Furthermore, the recently introduced TINNiK algorithm (Allman et al. 2024b) uses concordance factors computed from gene trees to construct the blobtree of networks with arbitrary level under the network multispecies coalescent model. Although such a blobtree still remains a tree, it does indicate in what areas of the underlying network reticulations may have occurred. It might also be worth looking for an extension of Squirrel to non-binary networks, where high-degree vertices are allowed which do not necessarily represent reticulate events.

In conclusion, Squirrel provides an efficient and combinatorially sound approach for reconstructing semidirected level-1 networks from dense sets of quarnets. The promising consistency scores achieved in our tests underscore Squirrel’s ability to retain network topology even when faced with noisy data. Together with our *δ*-heuristic, Squirrel allows rapid insight into large-scale sequence data. Looking forward, we hope that this approach can complement more time-intensive methods and support the preliminary exploration of network hypotheses.

## Materials and Methods

We start this section by presenting formal definitions surrounding phylogenetic networks and quarnets in the first subsection. The high level idea of Squirrel is described in the second subsection, while its subroutines are formalized in the third and fourth subsection. We end with the description of the *δ*-heuristic in the fifth subsection, and a brief description of the consistency and implementation of Squirrel in the sixth and seventh subsection, respectively.

### Phylogenetic networks and quarnets

#### Phylogenetic networks

A *rooted phylogenetic network* on a set of at least four leaves χ (representing a set of taxa) is a directed graph with a single root, no parallel edges and no directed cycles such that (i): the root has two children; (ii): each leaf (i.e. a vertex with no children) has one parent and is uniquely labeled by an element from χ ; (iii): all other vertices either have one parent and two children, or two parents and one child. A vertex of the latter type is a *reticulation (vertex)*, and the two edges directed towards it are *reticulation edges*. See Figure 9(a) for an example. *Semi-directed phylogenetic networks*, the type of network this paper is concerned with, can be obtained from a rooted phylogenetic network by suppressing its root and undirecting all edges except for the reticulation edges. For the sake of brevity, we refer to these networks simply as *semi-directed networks*. Since the reticulation edges remain directed, we can still refer to the reticulation vertices and edges of a semi-directed network (see Figure 9(b)). We call a semi-directed network *triangle-free* if it does not contain any triangles (3-cycles). Note that a semi-directed network without any reticulations is an (unrooted) phylogenetic tree in the usual sense.

**Figure 9:**
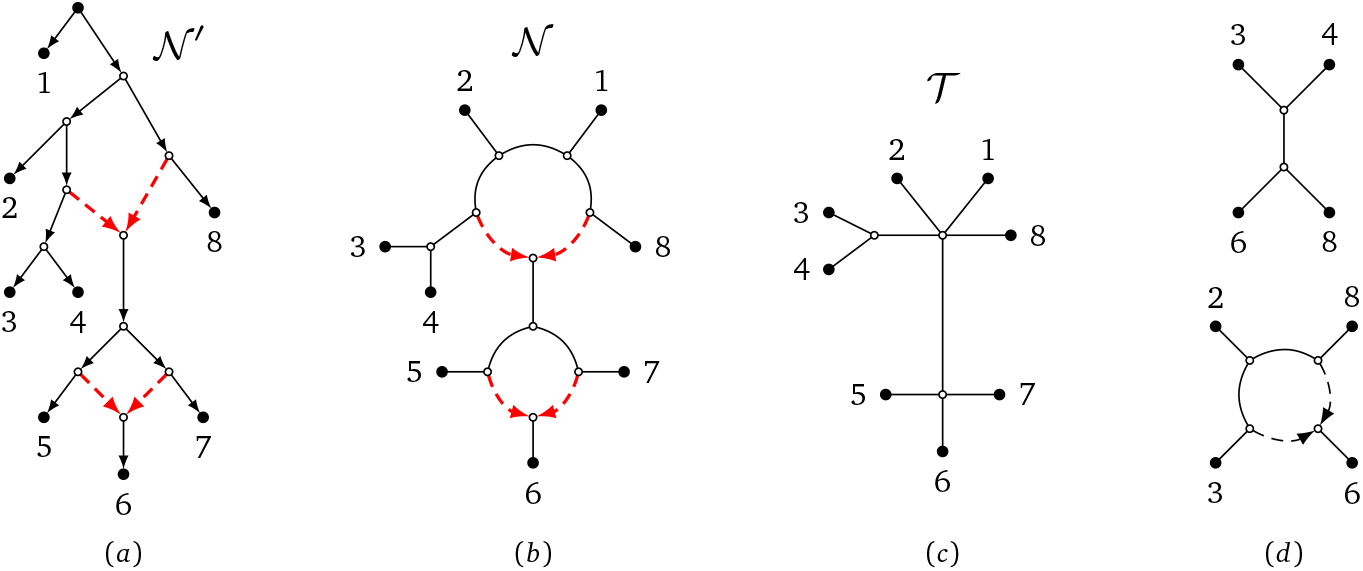
(a): A rooted phylogenetic level-1 network 𝒩 ^′^ on leaf set χ = {1, …, 8}, with the dashed reticulation edges pointing towards its reticulation vertices. (b): The triangle-free semi-directed level-1 network 𝒩 which can be obtained from 𝒩 ^′^ by suppressing its root and keeping only the dashed reticulation edges directed. (c): The blobtree 𝒯 of the semi-directed network 𝒩, obtained by collapsing all cycles into single vertices. (d): Two of the tf-quarnets induced by 𝒩. When ignoring the leaf labels, these are the two possible tf-quarnet shapes. The top tf-quarnet is a quartet tree and the bottom tf-quarnet is a 4-cycle.

In this paper, we consider semi-directed networks which are *level-1* (again see Figure 9(b)), meaning that every reticulation is part of exactly one undirected cycle (ignoring the directions of the reticulation edges). The (possibly non-binary) phylogenetic tree obtained by collapsing every such cycle into a single vertex is called the *blobtree* (or *tree of blobs*) of the semi-directed network (see Figure 9(c)).

Given a semi-directed network 𝒩 on χ, a partition *A B* of χ (with *A* and *B* both non-empty) is a *split* of if there exists an edge of whose removal disconnects the leaves in *A* from those in *B*. Such a split 𝒩 is *non-trivial* if the corresponding partition is non-trivial, that is, if |*A* |, |*B* | ≥ 2. As an example, 1, 2, 3, 4, 8} | {5, 6, 7} is a non-trivial split of the network from Figure 9(b). We sometimes omit the set notation for splits with few elements, meaning that we write *ab*|*cd* instead of the split {*a, b*}|{*c, d*} of the set {*a, b, c, d*}.

#### Quarnets

A semi-directed network *q* on a set of four leaves ℒ (*q*) = {*a, b, c, d*} is called a *(semi-directed) quarnet*. Recall that up to relabeling the leaves, there are six different level-1 quarnets (see Figure 1(c)). Here, we mostly focus on *tf-quarnets*: triangle-free level-1 quarnets. For a given leaf set ={*a, b, c, d*} and up to relabeling of the leaves, there are only two such tf-quarnets on : the *quartet tree* and the *4-cycle* (see Figure 9(d)). We often denote a quartet tree by its induced split (e.g. *ab cd*), while we describe a 4-cycle by its circular ordering (e.g. (*a, b, c, d*)) and mention the leaf below the reticulation separately. Note that tf-quarnets either have no non-trivial split at all, or they have exactly one non-trivial split (e.g. for χ = {*a, b, c, d*} the splits *ab*|*cd, ac*|*bd*, or *ad*|*bc*).

#### Squirrel: main algorithm

Squirrel uses as input a set 𝒬 of tf-quarnets on some leaf set χ with *n* = | χ | ≥ 4. In particular, this set needs to be *dense*, meaning that it contains exactly one tf-quarnet for each subset of four leaves of (see also the Introduction). Such a set can be created from a multiple sequence alignment using QNR-SVM (Barton et al. 2022), the MML algorithm (Martin et al. 2023) or our own *δ*-heuristic χ (see the fifth subsection of this section). We also allow for a function *w* : 𝒬 → [0, 1] to give weights to the tf-quarnets, which can e.g. be used to model confidence or bootstrap support. Unweighted tf-quarnets are assumed to have unit weights.

The main idea behind the Squirrel algorithm is to first build a sequence of *n* − 3 phylogenetic trees on the given *n* leaves, each one less refined than the other (see Algorithm 2). These trees will function as candidate blobtrees. By expanding all the high degree nodes in these trees into cycles (and introducing reticulations), we obtain a set of semi-directed candidate networks (see Algorithm 3). Finally, out of these networks, we choose the network 𝒩 with the highest *weighted tf-quarnet consistency score*, defined as

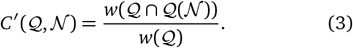

Here, 𝒬 is the input set of tf-quarnets and 𝒬 (𝒩) is the set of tf-quarnets which are induced by the output network 𝒩. A tf-quarnet *q* is *induced* by the network 𝒩 if it is the restriction of 𝒩 to ℒ (*q*), which is formally defined as the network obtained from 𝒩 by deleting all leaves not in ℒ (*q*) and exhaustively applying the following operations: deleting unlabeled leaves, deleting degree-2 reticulations, suppressing non-reticulate degree-2 vertices, suppressing parallel edges, and suppressing triangles. For completeness, we mention that an induced quarnet can be defined similarly but without suppressing the triangles.

The pseudo-code of Squirrel is shown as Algorithm 1. The blobtree construction algorithm (Algorithm 2) and the cycle expansion algorithm (Algorithm 3) are explained in detail in the following two subsections. Even though this is not specified in the pseudo-code, Squirrel does allow the user to specify an outgroup as input. Then, it makes sure that all candidate networks can be rooted using this outgroup (see also the fourth subsection of this section).

##### Algorithm 1

Squirrel

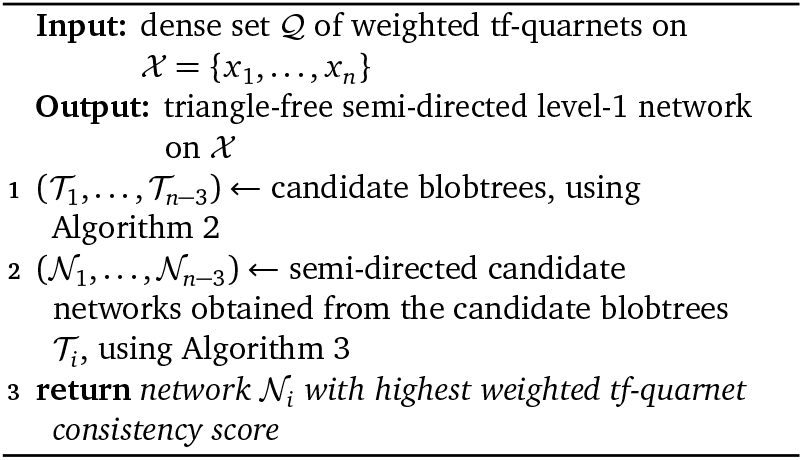

#### Squirrel: constructing candidate blobtrees

In the following three steps, we describe how Squirrel creates the sequence of candidate blobtrees on leaf set χ from the dense set 𝒬 of tf-quarnets. The pseudocode of this procedure is shown as Algorithm 2 at the end of this subsection.

**Step A1:** We first create a phylogenetic tree 𝒯 ^*^ on χ as described in Berry and Gascuel (2000). Their algorithm takes as input a (possibly non-dense) set 𝒬 ^′^ of quartet trees and returns as 𝒯 ^*^ the unique most refined phylogenetic tree on *χ* which does not induce a quartet with a different non-trivial split than one of the quartets in 𝒬 ^′^ (see Section A of the Supplementary Material for a more formal definition). By taking 𝒬 ^′^ to be the subset of quartet trees in our set of dense tf-quarnets (see line 1 of Algorithm 2) we can employ the algorithm from Berry and Gascuel (2000) to obtain 𝒯^*^ (see line 2 of Algorithm 2). As we show in Lemma A.2 of the Supplementary Material, in the case all tf-quarnets are induced by a unique network, 𝒯 ^*^ coincides with the blobtree of that network.

**Step A2:** Since the set 𝒬 (and thus 𝒬 ^′^) is constructed from real data, we expect there to be a fair amount of quartets that contradict each other. Hence, in practice, the tree 𝒯 ^*^ constructed in Step A1 will be highly unresolved. To remedy this problem, we use a method to refine the tree 𝒯 ^*^, specifically, an adapted version of the QuartetJoinING algorithm (Grünewald et al. 2009). QuartetJoining takes as input a function *ω* that assigns a non-negative real number to each possible non-trivial split of four leaves in *χ*. Starting with the star-tree with central vertex *v* and leaf set *χ*, QuartetJoining sequentially introduces edges between *v* and two of its neighbours (according to some criterion involving the function *ω*) until the tree is fully resolved.

In our case, we instead start with the tree 𝒯^*^ (which might already be partially resolved) and adapt QuarTETJoining to resolve *𝒯* ^*^ further. This eventually leads to a fully resolved phylogenetic tree 𝒯_1_ on χ, which functions as the first tree in our sequence of candidate blobtrees (see line 3 of Algorithm 2). In our adaptation, instead of considering all combinations of neighbours of the central vertex *v*, we consider all such combinations of neighbours of any of the internal (i.e. non-leaf) vertices with degree at least 4. We construct the function *ω* used as input to QuartetJoining as follows. For any tf-quarnet *q* ∈ *𝒬* with leaf set ℒ (*q*) = {*a, b, c, d*} such that *q* is a quartet tree (say with split *ab*| *cd*), we set *ω*(*ab* |*cd*) = *w*(*q*) for the input weight function *w* mentioned at the beginning of the previous subsection. All other non-trivial splits of four leaves of *χ* are assigned an *ω*-value of 0.

**Step A3:** Finally, we explain how we create the full sequence of candidate blobtrees from the phylogenetic tree 𝒯_1_. Given an edge *uv* of the tree 𝒯_1_ that induces a non-trivial split *A*| *B*, we collect all the quartet trees in 𝒬 for which their induced splits restrict to quartet splits of *A* |*B* in a set 𝒬^′^(*A*| *B*) by first defining 𝒬(*A* |*B*) = {*q* ∈ 𝒬 : *A* ∩ *ℒ*(*q*) = 2, *B* ∩ ℒ (*q*) = 2} and then 𝒬 ^′^(*A*| *B*) = {*q* ∈ (*A*| *B*) : *q* has split *A*∩ℒ (*q*) *B*∩ *ℒ* (*q*)}. This allows us to define the *split-support* of *uv* as

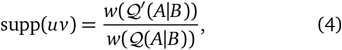

i.e. as the weighted ratio of the tf-quarnets in that support the split induced by the edge *uv*. For each of the *n* − 3 edges of the tree 𝒯_1_ we then compute this split-support (see line 4 of Algorithm 2). Afterwards, we sort the edges of 𝒯_1_ in increasing order, according to their split-support. To create the trees (𝒯 _2_, …, 𝒯_*n*− 3_), we keep contracting the least supported edge (see line 6 of Algorithm 2). In other words, the tree _*i*_ is obtained from 𝒯_1_ by contracting the *i* − 1 least supported edges. Crucial for our consistency proof in Section A of the Supplementary Material is that 𝒯_1_ is a refinement of 𝒯 ^*^, and therefore one of the trees in the sequence (𝒯_1_, …, 𝒯_*n*−3_) will be the tree 𝒯 ^*^.

##### Algorithm 2

Constructing candidate blobtrees

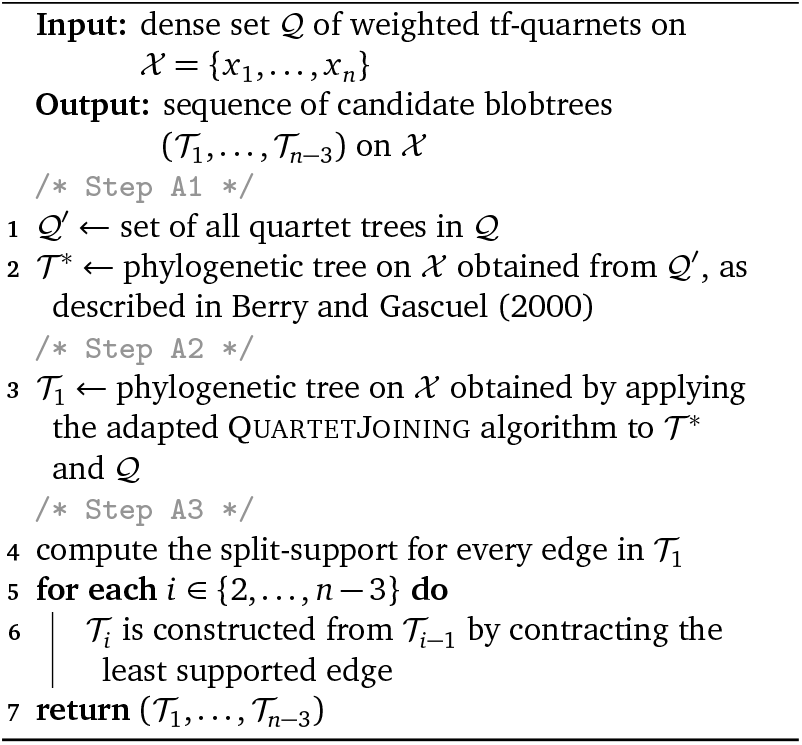

### Squirrel: expanding cycles in a tree

Once Squirrel has constructed the sequence of candidate blobtrees using Algorithm 2, we transform them into triangle-free semi-directed level-1 networks using the dense set of tf-quarnets 𝒬. In this subsection we describe how we transform a phylogenetic tree 𝒯representing one of our candidate blobtrees - into such a network. In particular, we replace every internal vertex of the given tree by a suitable cycle. Since our aim is to build triangle- free networks, we replace vertices incident to *s*≥ 4 edges by an *s*-cycle with a reticulation (see also the illustration in Figure 10). To this end, we repeat the following three steps for every such internal vertex *v* (starting with the ones with the highest degree). The corresponding highlevel pseudo-code is shown as Algorithm 3.

**Figure 10:**
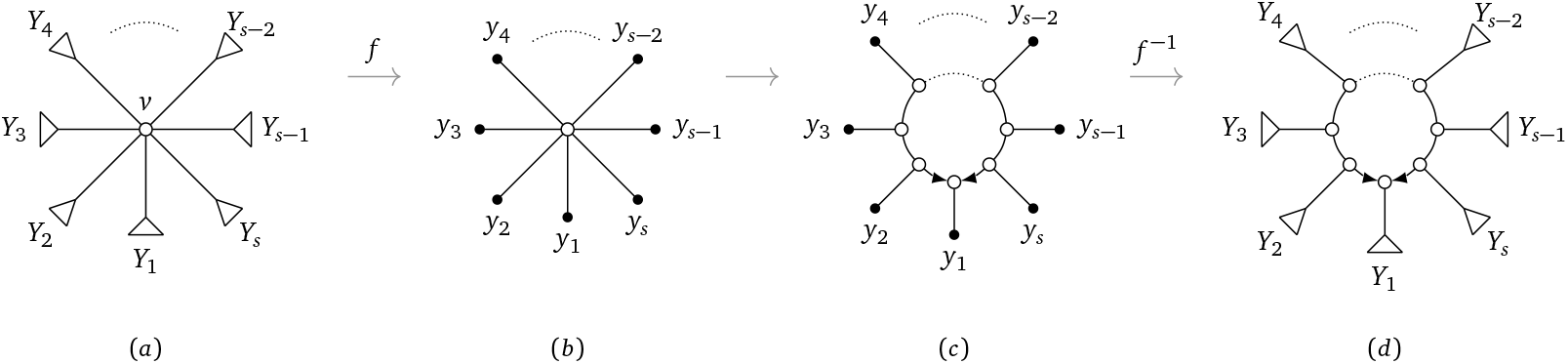
(a): A blobtree on some leaf set *χ* with an internal vertex *v* inducing the partition *Y*_1_ |… |*Y*_*s*_ of χ. (b): Illustration of the mapping *f* which maps every leaf *x* of χ to a leaf in {*y*_1_, …, *y*_*s*}_, depending on which set *Y*_*i*_ contains *x*. (c): Illustration of Step B2 and B3 of Squirrel, where the single internal vertex is replaced by a cycle. (d): Illustration of how the cycle on the leaves *y*_*i*_ is mapped back to a cycle on the sets *Y*_*i*_ with the inverse function *f* ^−1^.

**Step B1:** The first step in our approach is to assign a dense set of *representative tf-quarnets* 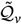 to each internal vertex *v* of 𝒯 with degree *s* ≥ 4. In particular, the set 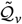 will be a dense set of tf-quarnets on the leaf set 𝒴 = {*y*_1_, …, *y*_*s*_}, where each *y*_*i*_ represents the set *Y*_*i*_ which is part of the partition *Y*_1_ |… |*Y*_*s*_ of χ induced by *v* (see Figure 10(a) and (b)). In the next step, these sets will then be used to determine by what cycle to replace *v*.

First, let *f* : χ → *𝒴* be the function that maps every leaf *x* ∈ χ, with *x* being in some set *Y*_*i*_, to the leaf *y*_*i*_ (see line 3 of Algorithm 3). To construct the tf-quarnets in 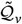 (see line 4 of Algorithm 3), we repeat the following procedure for every subset {*y*_*i*_, *y*_*j*_, *y*_*k*_, *y*_*l*_} of four leaves in *𝒴*. Let 𝒬 _*i,j,k,l*_ = {*q*∈ 𝒬 : ℒ (*q*) = {*x*_*i*_, *x* _*j*_, *x*_*k*_, *x*_*l*_} with *x*_*p*_ ∈ *Y*_*p*_ for all *p* ∈ {*i, j, k, l*} be the subset of 𝒬 containing only tf-quarnets with one leaf in each of the four sets *Y*_*i*_, *Y*_*j*_, *Y*_*k*_ and *Y*_*l*_. By relabeling the leaves of all tf-quarnets in 𝒬{_*i,j,k,l*}_ with the function *f*, we obtain a multiset of tf-quarnets which all have the same leaf set {*y*_*i*_, *y*_*j*_, *y*_*k*_, *y*_*l*_}. With slight abuse of notation, we denote this multiset by *f* (𝒬_{*i,j,k,l*}_). Then, we choose one of the tf-quarnets in the multiset *f* (𝒬{_*i,j,k,l*}_) to assign to 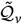 as the tf-quarnet on the four-leaf set {*y*_*i*_, *y*_*j*_, *y*_*k*_, *y*_*l*_} (see next paragraph). As mentioned before, this is repeated for every subset {*y*_*i*_, *y*_*j*_, *y*_*k*_, *y*_*l*_} of four leaves in 𝒴, resulting in a dense set of tf-quarnets on *𝒴*.

To choose a tf-quarnet from the multiset *f* (𝒬 _{*i,j,k,l*}_), we first choose its *skeleton*: its underlying undirected graph. In particular, for each of the six possible skeletons *t* (three quartet trees and three undirected 4-cycles) we let *w*(*t*) be the sum of weights of all tf-quarnets in *f* (𝒬 {_*i,j,k,l*}_) with the given skeleton *t*. We then choose the skeleton *t* with the highest weight (with ties resolved randomly) and assign it a new weight of *w*(*t*)*/w*(*f* (𝒬 {_*i,j,k,l*}_)). Note that in the unweighted case this simply means that we choose the skeleton that appears most in the multiset. We first choose the skeleton since determining the location of the reticulation in a quarnet from data seems especially hard (Martin et al. 2023). If our chosen skeleton is one of the quartet trees, we assign that as our tf-quarnet on {*y*_*i*_, *y*_*j*_, *y*_*k*_, *y*_*l*_}. On the other hand, if one of the undirected 4-cycles appears most, we still need to determine the location of the reticulation. This is done by checking which leaf appears most often below the reticulation in all 4-cycles with the chosen skeleton.

As an example of this voting procedure to choose a tf-quarnet from the multiset *f* (𝒬 _*i,j,k,l*_), suppose our multiset *f* 𝒬 _{*i,j,k,l*}_contains only tf-quarnets with weight 1 and is as in Figure 11.

**Figure 11:**
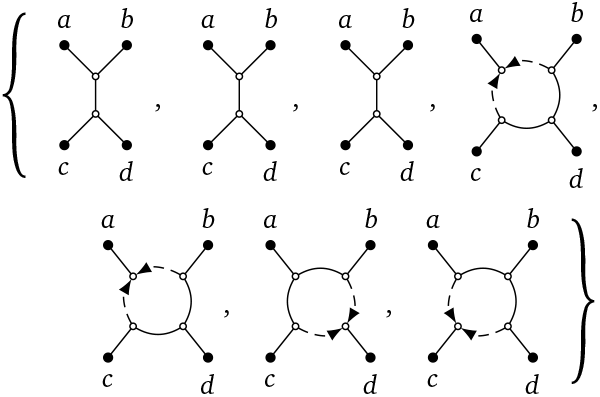
A multiset of 7 tf-quarnets on leaf set {*a, b, c, d*}.

Then, we choose the 4-cycle with circular ordering (*a, b, c, d*) as our skeleton, after which we assign *a* to be the leaf below the reticulation. The new tf-quarnet is then a 4-cycle with a weight of 4*/*7 because that 4-cycle appears 4 times out of a total of 7 tf-quarnets.

**Step B2:** The next step of our approach is to determine a circular ordering of the leaves in the set *𝒴* based on the tf-quarnets in 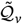.Note that we repeat this for every internal vertex *v* of 𝒯 with degree at least 4. First, we use the set 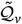 to create a distance 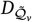 between every pair of leaves in *𝒴* (see line 5 of Algorithm 3). Formally, given two leaves *a* and *b* in 𝒴, we define the distance 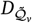 as follows:

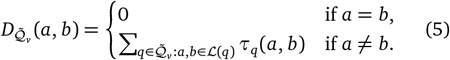

For every tf-quarnet 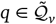 the exact value of *τ* depends on the weight of *q* and the position of the leaves *a* and *b* within it. In particular, the values are defined on the skeleton of the tf-quarnets and hence do not depend on the position of the reticulations. Given two leaves *a* and *b* of a tf-quarnet *q* (with weight *w*(*q*) ∈ [0, 1]), we define *τ*_*q*_ as follows:

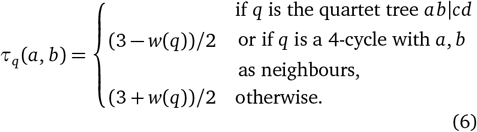

Here, we say that two leaves of a 4-cycle are *neighbours* if they are not on opposite sides of the cycle. The *τ*_*q*_ -values reduce to 1 or 2 for tf-quarnets *q* with a weight of 1. Specifically, two leaves on the same side of a split in a quartet tree *q* have a *τ*_*q*_ -value of 1, otherwise they have a *τ*_*q*_ -value of 2. Similarly, two neighbouring leaves in a 4-cycle *q* have a *τ*_*q*_ -value of 1, while two opposite leaves have a *τ*_*q*_ -value of 2. See Figure 12 for an illustration of these values. Note that these pairwise distances between leaves resemble the *quartet distances* used in NANUQ (Allman et al. 2019) and NANUQ^+^ (Allman et al. 2024a).

**Figure 12:**
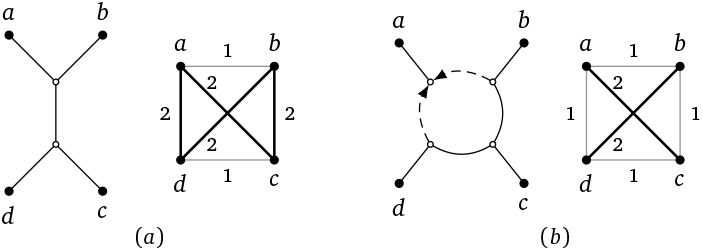
Two tf-quarnets *q* with leaf set {*a, b, c, d* : a} quartet tree (a) and 4-cycle (b). The values *τ*_*q*_ (as defined by eq. (6), assuming the quarnets have weight 1) between any two leaves are illustrated by the two complete graphs, where the thin grey edges have length 1 and the thick black length 2.

Once the distances 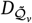 are computed, we create a complete graph *G* with vertex set *𝒴*, where the distances between the vertices are given by 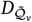.By solving the Trav-ELLING Salesman Problem (TSP) on this graph we obtain a circular ordering of the elements 𝒴 in (see line 6 of Algorithm 3). The goal of a TSP instance is to find a shortest *Hamiltonian cycle* (or *TSP-tour*): a cycle that visits each vertex exactly once. The default setting for Squirrel is to use the Held-Karp algorithm (Bellman 1962; Held and Karp 1962) for up to and including 13 leaves and to use simulated annealing to heuristically solve instances with more leaves. To obtain true consistency (see Section A of the Supplementary Material) this setting can be changed to always solve TSP to optimality, at the cost of a longer running time.

**Step B3:** After solving TSP, Squirrel obtains a circular ordering *θ* of the leaves in *𝒴*. It remains to determine which leaf *y*_*i*_ needs to be the leaf below the reticulation in the resulting cycle. To ensure Squirrel always returns a valid (that is, rootable) semi-directed network, we create a *reticulation ranking ρ* of the leaves in 𝒴 instead of picking a single leaf (see line 7 of Algorithm 3). If the set 𝒴 contains at least five elements, we order them according to how often they appear in a 4-cycle of 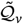 (as defined in Step B1). That is, the first leaf in our ranking *ρ* appears most often in a 4-cycle and is our first option to be the leaf below the reticulation. The case where |𝒴 |= 4 is special, since 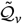 then only contains a single quarnet. If this is a 4-cycle, then the leaf below the reticulation of that 4-cycle is the first leaf in our ranking *ρ*. The other three leaves (or in the case that the single tf-quarnet is a quartet tree: all four leaves) are ordered randomly.

Finally, we map every leaf *y*_*i*_ back to the corresponding leaf set *Y*_*i*_ of the original tree *𝒯* with the inverse function *f* ^−1^. While slightly abusing notation, this results in an ordering *f* ^−1^(*θ*) of the sets *Y*_*i*_. Then, we replace the internal vertex *v* in the tree *𝒯* by a cycle that follows this ordering *f* ^−1^(*θ*) (see line 9 of Algorithm 3 and Figure 10(b) and (c) for an illustration). We determine the location of the reticulation by looking at the first element *ρ*_1_ of the reticulation ranking *ρ*. In particular, we let the leaf set in *f* ^−1^(*ρ*_1_) be below the reticulation (again see line 9 of Algorithm 3). This could possibly create a partially constructed network that is *invalid*: one without a valid root location (e.g. if two reticulations are oriented towards each other). Hence, if this is the case we instead pick the leaves in *f* ^−1^(*ρ*_2_). If this is still an invalid option, we keep iterating through the ranking *ρ* until we find a valid partial network (see line 10 of Algorithm 3). Note that This procedure ensures that we always return a valid semi-directed network at the end of Algorithm 3. Our implementation of Squirrel also allows the user to specify a known outgroup. Then, a (partially constructed) semi-directed network is only valid if it is not only rootable, but if it can also be rooted at the edge incident to the outgroup. Iterating through the reticulation ranking ensures that we always return a valid semi-directed network at the end of Algorithm 3, even in the case of a specified outgroup (see Lemma A.6 in Section A of the Supplementary Material for a proof).

### *δ*-heuristic: inferring quarnets from sequence data

As explained before, two model-based methods that use algebraic invariants exist to generate tf-quarnets (Barton et al. 2022; Martin et al. 2023). To allow Squirrel to function as a stand-alone tool, we also include a method to infer weighted tf-quarnets from a multiple sequence alignment (MSA) on a set of taxa χ: the *δ*-heuristic. Our *δ*-heuristic is based on the concept of *δ*-plots, which function as a measure of treelikeness for sets of four taxa and which were able to pick out recombinants in many simulations (Holland et al. 2002). The algorithm also resembles some aspects of the heuristic to generate trinets from sequences in Oldman et al. (2016). We are now ready to present the steps to create a dense set of weighted tf-quarnets from an MSA on leaf set *χ*.

#### Algorithm 3

Expanding cycles in a tree

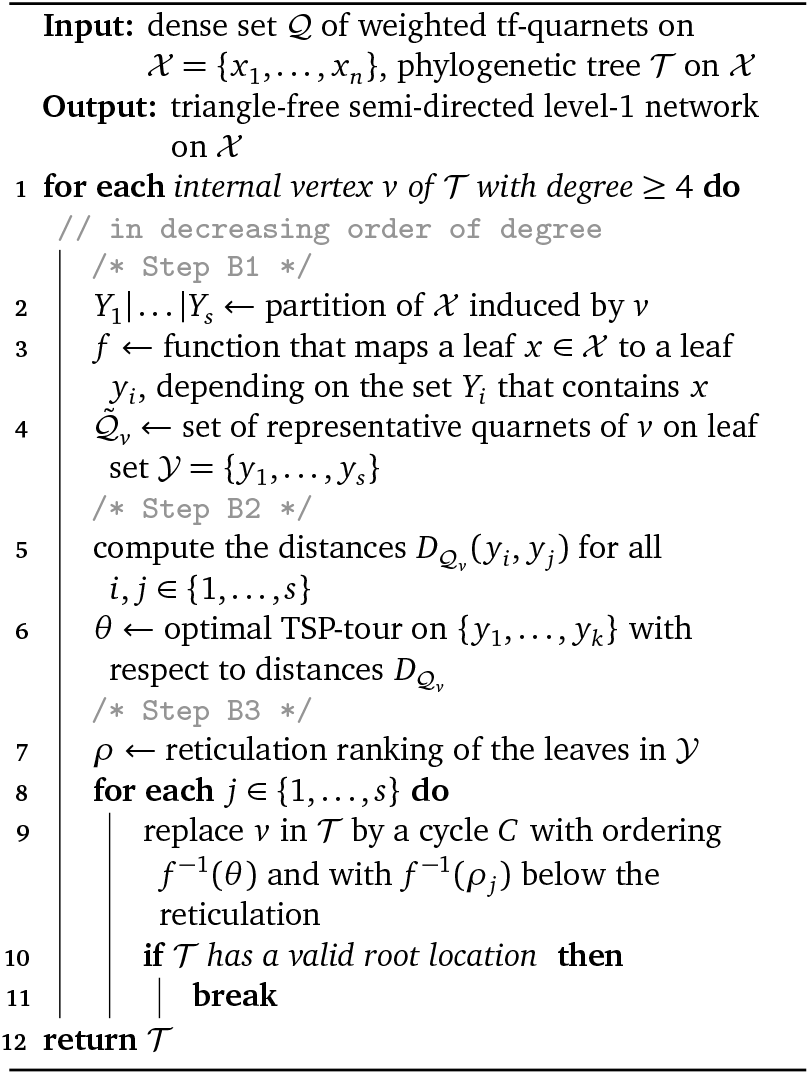

**Step I:** For each pair of taxa {*a, b*} we consider the gapfree subalignment of the MSA on {*a, b*}. That is, we consider only the columns where both taxon *a* and *b* contain no gaps. Using this subalignment, we assign a distance value *h*_*ab*_ to the pair {*a, b*}. In particular, *h*_*ab*_ is the *normalized Hamming distance*: the number of columns of the subalignment where taxon *a* and *b* differ, divided by the total length of the subalignment. Recall that if a tf-quarnet on {*a, b, c, d*} has a non-trivial split, it has one of the three splits *ab*| *cd, ac*| *bd* or *ad*| *bc*. For each four taxon subset and for each of these three splits, say *ab* | *cd*, we then let *h*_*ab*|*cd*_ = *h*_*ab*_ + *h*_*cd*_.

The *δ*-value (introduced in Holland et al. 2002) of such a subset {*a, b, c, d*} of *χ* is now defined as follows (assuming we have that *h*_*ab*|*cd*_ ≥ *h*_*ac*|*bd*_ ≥ *h*_*ad*|*bc*_):

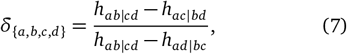

where *δ*_{*a,b,c,d*}_ = 0 if *h*_*ab*|*cd*_ = *h*_*ac*|*bd*_ = *h*_*ac*|*bd*_. Intuitively, the *δ*-value indicates how much support there is from the subalignment that the tf-quarnet on {*a, b, c, d*} has a split. That is, if the value of *δ*_{*a,b,c,d*}_ is close to 1, we expect the split *ab*|*cd* to be present.

**Step II:** With the *δ*-values computed for each subset of four taxa, we partition the 4-taxa sets into two subsets *S*_*λ*_ and *F*_*λ*_ for a predefined threshold value *λ* ∈ (0, 1). The set *S*_*λ*_ will contain all 4-taxa subsets for which the *δ*-value is at least *λ*, while the set *F*_*λ*_ contains those sets with an *δ*-value smaller than *λ*. We then expect the sets in *S*_*λ*_ to come from a tf-quarnet with a non-trivial split, while those in *F*_*λ*_ are likely to have come from 4-cycle tf-quarnets. Experiments from Holland et al. (2002) show that an average *δ*-value higher than 0.3 is often enough to determine whether recombination was present (or equivalently, whether a tf-quarnet has a non-trivial split). Hence, we settle for a value of *λ* = 0.3.

**Step III:** Every 4-taxa set {*a, b, c, d*} in *S*_*λ*_ is assigned a quartet tree. Its split is simply determined by the split *s* ∈ {*ab*| *cd, ac* |*bd, ad*| *bc*} for which *h*_*s*_ is the highest. On the other hand, the sets in *F*_*λ*_ will be assigned a 4-cycle. Observe that any 4-cycle tf-quarnet with circular ordering (*a, c, b, d*) (irrespective of the position of the reticulation) can be turned into the quartet trees with splits *ac* |*bd* or *ad*| *bc* by deleting exactly one reticulation edge, while this is not possible for the quartet tree with split *ab*| *cd*. Assuming that the taxa set {*a, b, c, d*} is in the set *F*_*λ*_ and that *h*_*ac*|*bd*_ *>*≥ *h*_*ac*|*bd*_ ≥ *h*_*ac*|*bd*_, therefore assign a 4-cycle with circular ordering (*a, c, b, d*) to the taxa set. This aligns with the group-based models (see e.g. Gross et al. 2021; Barton et al. 2022) which also assume that DNA independently evolves along the trees that can be obtained from a network by deleting reticulation edges.

We also assign a weight *w*(*q*) to each tf-quarnet *q*, corresponding to the difference its *δ*-value has from *λ*. In some sense, this weight signifies the confidence we have in having estimated the correct tf-quarnet. In particular,

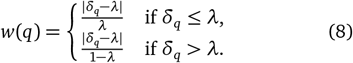

**Step IV:** It remains to determine where to place the reticulations in the 4-cycles obtained from the set *F*_*λ*_. Taking inspiration from Holland et al. (2002) and Oldman et al. (2016), we first compute the value *δ*(*x*) for each taxon *x*, defined as the mean value of all *δ*-values for four-taxon sets containing *x*. For each 4-cycle, we then let the leaf *x* with the highest *δ*(*x*)-value be below the reticulation.

### Consistency of Squirrel

In Section A of the Supplementary Material we prove that Squirrel is combinatorially consistent given an unweighted dense set of tf-quarnets. We use the word ‘combinatorially’ to emphasize that we do not make any claims regarding statistical consistency. More formally, we prove the following theorem.

#### Theorem 1.

*Let 𝒩 be a triangle-free semi-directed level-1 network and let* 𝒬 *be the set of unweighted tf-quarnets induced by 𝒩, then* Squirrel *applied to* 𝒬 *reconstructs 𝒩*.

The first ingredient of the proof is the fact that if a set of tf-quarnets is induced by a network, the tree 𝒯^*^ is equal to the blobtree of that network. The other important step of the proof is to show that in this case the distances *D* (as defined in eq. (6)) form a *Kalmanson metric* (Kalmanson 1975), which have nice properties with respect to the Travelling Salesman Problem.

### Implementation

A graphical user interface (implemented in Python) of Squirrel and the *δ*-heuristic is freely available at https://github.com/nholtgrefe/squirrel. The program takes as input a sequence alignment in NEXUS or FASTA format, or a file specifying a dense set of tf-quarnets (e.g. coming from QNR-SVM (Barton et al. 2022) or the MML algorithm (Martin et al. 2023)). The interface allows the user to specify an optional outgroup, view the different generated candidate networks, and export them in the eNewick file-format (Cardona et al. 2008) (with an arbitrary rooting if no outgroup was specified).

## Supplementary Material

Supplementary material is available at Molecular Biology and Evolution online (http://www.mbe.oxfordjournals.org/).

## Data availability statement

The generated networks, Python scripts, sequence alignments and numerical results of the experiments in this paper are available at https://github.com/nholtgrefe/squirrel.

## Acknowledgements

This work received funding from grants OCENW.M.21.306 (NH, LvI & MJ) and OCENW.KLEIN.125 (LvI & MJ) of the Netherlands Organisation for Scientific Research (NWO). Part of this work was done while some of the authors were in residence at the Institute for Computational and Experimental Research in Mathematics (ICERM) in Providence (RI, USA) during the *Theory, Methods, and Applications of Quantitative Phylogenomics* program (supported by grant DMS-1929284 of the National Science Foundation (NSF)). We thank the authors of Allman et al. (2024a) for making us aware of their new approach and the reviewers for their helpful comments and suggestions to improve the paper.

## A Proof of consistency

In this section we prove that Squirrel is combinatorially consistent given an unweighted dense set of tf-quarnets:

### Theorem 1.

*Let* 𝒩 *be a triangle-free semi-directed level-1 network and let* 𝒬 *be the set of unweighted tf-quarnets induced by* 𝒩, *then* Squirrel *applied to* 𝒬 *reconstructs* 𝒩.

We start by giving a formal definition (taken from Berry and Gascuel (2000), originally by Bandelt and Dress (1986)) of the tree 𝒯 ^*^ constructed in Section 4.3, after which we prove that it is a combinatorially consistent estimator of the blobtree of the network 𝒩. ^′^

Given a non-trivial split *A*|*B* of *χ*, we let 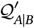 be the set of all possible quartets that *agree* with the split *A*|*B*. That is, for every *a*_1_, *a*_2_ ∈ *A* and 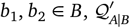 contains the quartet *q* with split *a*_1_*a*_2_|*b*_1_ *b*_2_. Given a (possibly non-dense) set of quartets 𝒬^′^ on χ, we let *S*^*^ be the maximal set of splits of χ such that 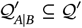 for all splits *A*|*B* in *S*^*^. In other words, *S*^*^ is the maximal set of splits such that 𝒬^′^ contains all quartets that agree with these splits. Lastly, we define the set 𝒬^*^ as the subset of 𝒬^′^ that contains exactly these quartets, i.e. 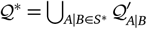. In Berry and Gascuel (2000) it is shown that the set 𝒬^*^ can also be characterized as the unique maximum subset of 𝒬^′^ that is *tree-like*, meaning that there exists a phylogenetic tree 𝒯 on χ with 𝒬^*^ as the set of resolved quartets (i.e. the quartets contain a non-trivial split) it induces. This is the tree 𝒯^*^(𝒬 ^′^) (or simply 𝒯^*^ if the set 𝒬^′^ is clear). Note that this tree is unique since a phylogenetic tree is uniquely determined by its quartets (Colonius and Schulze 1981), and thus by its resolved quartets.

We are now ready to prove that the tree 𝒯 ^*^ constructed by Squirrel is exactly the blobtree of 𝒩, provided the tf-quarnets induced by 𝒩 are used as input. Recall that to construct the tree 𝒯 ^*^ given a dense set 𝒬 of tf-quarnets, Squirrel first creates a set of quartets 𝒬^′^ ⊆ 𝒬 by throwing out all the 4-cycles. This set can then be used by to obtain the tree 𝒯 ^*^ as described in Berry and Gascuel (2000).

### Lemma A.2.

*Let* 𝒩 *be a triangle-free semi-directed level-1 network, let* 𝒬 *be the set of tf-quarnets induced by* 𝒩 *and let* 𝒬^′^ ⊆ 𝒬 *be the set of quartet trees in* 𝒬. *Then, the blobtree of* 𝒩 *is equal to the tree* 𝒯 ^*^(𝒬^′^).

*Proof*. Let 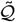 be the set of resolved quartets (i.e. quartets with a non-trivial split) induced by the blobtree 𝒯 of 𝒩. It is enough to show that 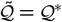 since phylogenetic trees are uniquely determined by their induced resolved quartets (Colonius and Schulze 1981) and, by definition, 𝒬^*^ is the set of induced resolved quartets of 𝒯 ^*^((𝒬^′^).

Let 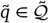 be a resolved quartet with split 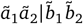.This means that there is some non-trivial split *A* |*B* in 𝒯 (and thus in 𝒩) with 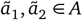 and 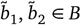.Now let *a*_1_, *a*_2_ ∈ *A* and *b*_1_, *b*_2_ ∈ *B* be arbitrary. By Theorem 5.1 in Huber et al. (2024) the quarnet *q* with *ℒ* (*q*) = {*a*_1_, *a*_2_, *b*_1_, *b*_2_} that is induced by 𝒩 then has split *a*_1_*a*_2_|*b*_1_ *b*_2_. This directly implies that there is a quartet in 𝒬^′^ with split *a*_1_*a*_2_|*b*_1_ *b*_2_. Consequently, 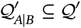 and 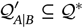 Because 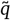 agrees with *A*|*B* we have 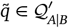,so we obtain 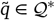.Since 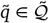 was arbitrary, 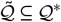.

Now let *q*^*^ ∈ 𝒬^*^ be an arbitrary resolved quartet with split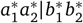.By definition, *q*^*^ must be in 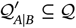 for some non-trivial split *A*|*B* (with 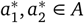 and 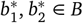). Since 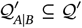,for any *a*_1_, *a*_2_ ∈ *A* and *b*_1_, *b*_2_ ∈ *B* the quartet *q* with split *a*_1_*a*_2_ | *b*_1_ *b*_2_ is in 𝒬^′^. By (Huber et al. 2024, Thm. 5.1), this means that *A B* is a split in 𝒩 and thus in 𝒯. But *q*^*^ is a resolved quartet of 𝒯, so 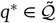,Since *q*^*^ was arbitrary, this shows that 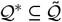.

We continue with proving that the approach in Step B2 of Squirrel correctly determines the ordering of each cycle. For the rest of this section, we consider to be a dense set 𝒬 of unweighted tf-quarnets induced by a *sunlet network*: a semi-directed level-1 network consisting of a single cycle with pendent leaves. This will then be enough to prove the consistency at the end of this section.

Clearly, a sunlet network induces a circular ordering of its leaf set 𝒴. We now create an explicit formula for the distance function *D*_𝒬_ defined in Section 4.4, assuming the set 𝒬 contains tf-quarnets induced by a sunlet network.

Note that a distance function *D* on a finite set of elements 𝒴 is a *metric* if (i) it is symmetric; (ii) the triangle inequality holds; (iii) the distance between two elements is zero if and only if the elements are equal.

### Lemma A.3.

*Let* 𝒩 *be a sunlet network on* 𝒴 = {*y*_1_, …, *y*_*n*_}, *let θ* = (*y*_1_, …, *y*_*n*_) *be a circular ordering of* 𝒴 *induced by* 𝒩 *such that y*_1_ *is the leaf below the reticulation. If* 𝒬 *is the set of unweighted tf-quarnets induced by* 𝒩, *then D*_𝒬_ *is a metric on* 𝒴 *and it can be expressed as*

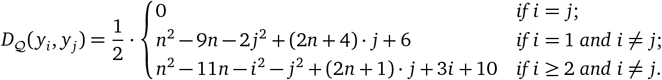

*Proof*. Clearly, *D*_𝒬_ is symmetric and two leaves have distance zero if and only if they are the same. To see that the triangle inequality holds, note that *D*_𝒬_ is defined as a sum of *τ*-values. Since those values all adhere to the triangle inequality (assuming the tf-quarnets all have weight 1), it readily follows that *D*_𝒬_ also has the same property. Hence, it is a metric.

We will now find the expression for the distances defined by *D*_𝒬_. Given two leaves *y*_*i*_ and *y*_*j*_ with 1 ≤ *i < j* ≤ *n*, let *r*_*i*_ = *i* −2 be the number of leaves between *y*_1_ and *y*_*i*_, and let *t* _*j*_ = *n* −*j* be the number of leaves between *y*_*j*_ and *y*_1_ (on the side of the cycle containing leaf *y*_*n*_). Lastly, let *s*_*i j*_ = *j*− *i* −1 be the number of leaves between *y*_*i*_ and *y*_*j*_. See Figure A.13 for an illustration.

Now let *i* = 1 and le t *i < j* ≤ *n* be arbitrary. Any tf-quarnet in 𝒬 containing both *y*_*i*_ and *y*_*j*_ will be a 4-cycle. In particular, in 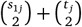 of these 4-cycles, *y*_*i*_ and *y*_*j*_ will be opposite leaves. On the other hand, in *s*_1*j*_ · *t* _*j*_ of these 4-cycles they will be neighbours.

We now let 1 *< i < j*≤ *n* be arbitrary. In this case, any tf-quarnet in 𝒬 containing both *y*_*i*_ and *y*_*j*_ will either be a quartet tree or a 4-cycle. All 4-cycle tf-quarnets must contain leaf *y*_1_ as well. Then, in *r*_*i*_ + *t* _*j*_ of them *y*_*i*_ and *y*_*j*_ are opposite leaves, while in *s*_*i j*_ of them they are neighbours. All quartet trees that include *y*_*i*_ and *y*_*j*_, do not include leaf *y*_1_. In 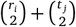 of the quartet trees, *y*_*i*_ and *y*_*j*_ are on the same side of the split, i.e. there is no ‘*y*_*i*_| *y*_*j*_ split’. The number of quartet trees that do have a ‘*y*_*i*_| *y*_*j*_ split’ is 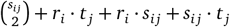. Filling in the distances defined by *D*_𝒬,_, we thus obtain for each 1 ≤ *i < j* ≤ *n*:

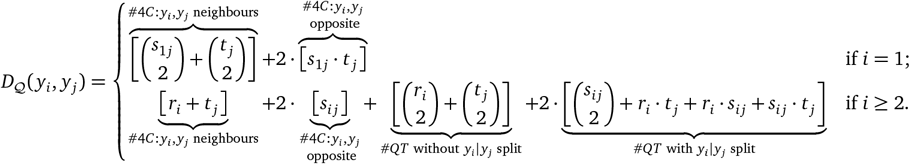

If we fill in the formulas for *r*_*i*_, *t* _*j*_ and *s*_*i j*_, we obtain

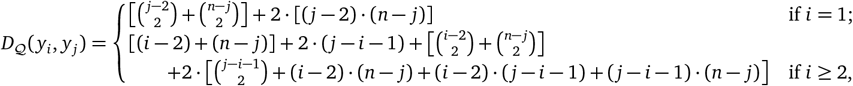

which reduces to

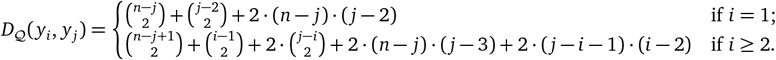

After writing out the binomial coefficients and expanding all brackets, one obtains the desired formula for those *y*_*i*_ and *y*_*j*_ with 1 ≤ *i < j* ≤ *n*. The other cases follow since *D*_𝒬_ is a metric.

**Figure A.13:**
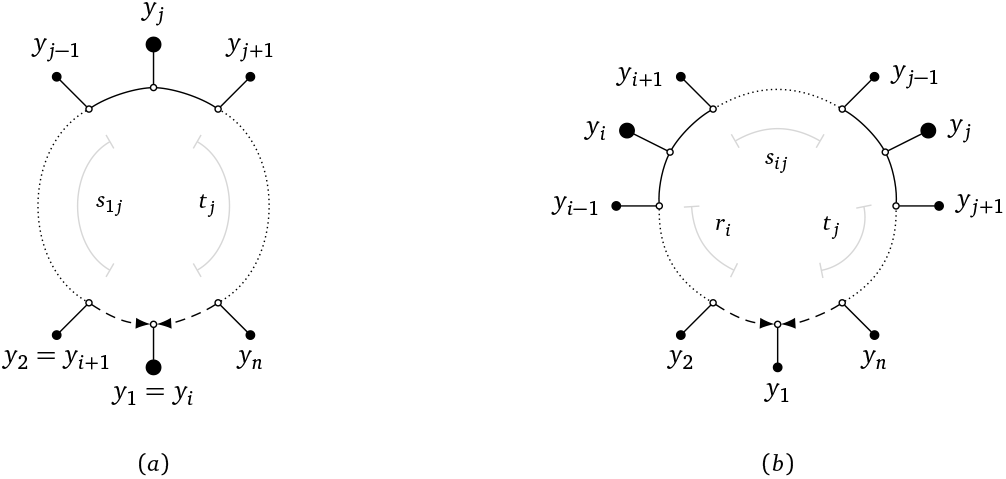
Illustration of the definitions of *r*_*i*_, *s*_*i j*_ and *t* _*j*_ in the proof of Lemma A.3. Subfigure (a) is the case where *i* = 1 and subfigure (b) is the case where *i >* 1.

The explicit formula derived in the previous lemma allows us to prove that *D*_𝒬_ is *Kalmanson* (Kalmanson 1975) with respect to the circular ordering *θ* of the leaves. Such a metric *D* defined on a set of elements 𝒴 has the following nice property: it is easy to find an optimal TSP-tour in the complete graph 𝒴 on with distances defined by *D*. Moreover, the corresponding ordering *θ* defines an optimal TSP-tour. Formally, these metrics are defined as follows.

### Definition A.4

(Kalmanson metric). Let *θ* = (*y*_1_, …, *y*_*n*_) be an ordering of a finite set of elements *𝒴* = {*y*_1_, …, *y*_*n*_}. A metric *D* on *𝒴* is *Kalmanson* with respect to *θ* if both of the following conditions hold:

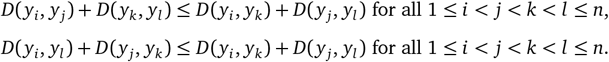

In the next lemma we prove that *D*_𝒬_ is a Kalmanson metric with respect to a circular ordering of the leaves induced by a sunlet network. From this it follows that TSP can be used to find such an ordering.

### Lemma A.5.

*Let* 𝒩 *be a sunlet network on* 𝒴 = {*y*_1_, …, *y*_*n*_} *and let* 𝒬 *be the set of unweighted tf-quarnets induced by* 𝒩. *Then, a circular ordering θ of* 𝒴 *is induced by* 𝒩 *if and only if θ is an optimal TSP-tour defined by D*_𝒬_.

*Proof*. Let *θ* = (*y*_1_, …, *y*_*n*_) be an ordering of 𝒴 induced by 𝒩. We will first show that *D*_𝒬_ is Kalmanson with respect to *θ*. Then, it will follow from Kalmanson (1975) that *θ* is an optimal TSP-tour. Without loss of generality, we can assume that *y*_1_ is the leaf below the reticulation since being Kalmanson is invariant under cyclic permutations (see e.g. De?ineko et al. 1997). We will now prove that *D*_𝒬_ is Kalmanson with respect to *θ* by checking the Kalmanson conditions from Definition A.4. For easier notation, we multiply the expressions by 2. Now let 1 ≤*i < j < k < l* ≤*n* be arbitrary. Using the explicit formula of Lemma A.3, we then distinguish between the cases where *i* = 1 and *i >* 1. Note that the *n*^2^ −9*n* + 6 and *n*^2^ −11*n* + 10 parts cancel out in all conditions.

*Case 1: i* = 1. To prove the first condition we use that *j < k*:

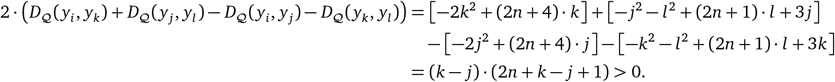

For the second condition, we use the fact that 3 *< k < l*:

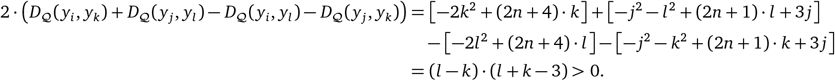

*Case 2: i >* 1. The first condition follows from the fact that *k > j* and 2*n >* 2:

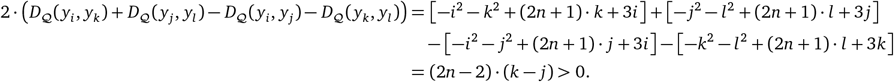

The second condition is trivial:

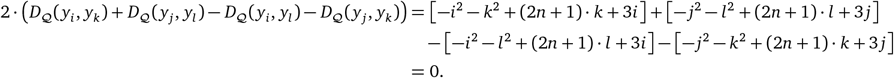

It remains to show that any circular ordering *φ* of 𝒴 that is not induced by 𝒩 is not an optimal TSP-tour. To see this, we argue that the total TSP-distance of any such ordering *φ* can always be decreased by swapping two specific adjacent leaves. In particular, any such ordering *φ* will have four leaves {*y*_*i*_, *y*_*j*_, *y*_*k*_, *y*_*l*_} (with 1≤ *i < j < k < l*≤ *n*) adjacent in the ordering *φ* but ordered as (*y*_*i*_, *y*_*k*_, *y*_*j*_, *y*_*l*_) (if *i* ≠ 1), or ordered as (*y*_1_, *y*_*k*_, *y*_*j*_, *y*_*l*_) or (*y*_*j*_, *y*_1_, *y*_*k*_, *y*_*l*_) (if *i* = 1). In all three cases, we can swap two of these adjacent leaves to decrease the total distance, since the corresponding three Kalmanson inequalities are strict inequalities.

We are now ready to present the proof of Theorem 1.

*Proof of Theorem 1*. Since the set of tf-quarnets 𝒬 is induced by the network 𝒩, we know by Lemma A.2 that the tree 𝒯 ^*^ (as constructed in Step A1) is equal to the blobtree of 𝒩. The tree 𝒯_1_ (in Step A2) is a *refinement* of 𝒯 ^*^. That is, 𝒯 ^*^ can be obtained from 𝒯_1_ by contracting edges. Recall that the tree 𝒯 ^*^ is the unique most refined tree on *χ*such that no tf-quarnet in 𝒬 contradicts a split of 𝒯 ^*^. This means that if we keep contracting the least supported split of 𝒯_1_ (see equation (4) in Step A3 of Section 4.3), we eventually end up with 𝒯 ^*^. Therefore, the tree 𝒯 ^*^ (and thus the blobtree of 𝒩) is part of the sequence (𝒯_1_, …, 𝒯_*n*−3_).

Next, we show that given the blobtree 𝒯 ^*^ of 𝒩 (and assuming the set 𝒬 is induced by 𝒩), Step B of Squirrel correctly constructs the network 𝒩. We first show this is true when 𝒩 is a sunlet network (and so 𝒯^*^ is an unresolved tree with a single internal vertex). Then, the set 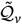 constructed in Step B1 is the same as the original set 𝒬of tf-quarnets (up to the relabeling defined by *f*). From Lemma A.5 we then know that the optimal TSP-tour using the distances 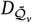 will correspond to the circular ordering induced by 𝒩. Whenever the sunlet network 𝒩 has at least five leaves, only the leaf below the reticulation will appear in every 4-cycle tf-quarnet induced by 𝒩. On the other hand, if 𝒩 has only four leaves it only induces one tf-quarnet: a 4-cycle with the correct leaf below its reticulation. Hence, Squirrel correctly picks the reticulation vertex in Step B3. Thus, in the case that 𝒩 is a sunlet network Squirrel constructs 𝒩 from 𝒯 ^*^.

Whenever 𝒩 is not a sunlet network, the proof is similar. To be precise, for every internal vertex *v* with induced partition *Y*_1_| … |*Y*_*s*_, let 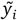 be an arbitrary leaf of *Y*_*i*_. Then, up to the relabeling defined by *f*, the set 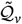 (as constructed in Step B1) is equal to the set of tf-quarnets induced by the sunlet network 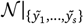 (the restriction of 𝒩 to 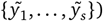.

Using a similar argument as above, Step B3 then correctly reconstructs the cycle in the sunlet network 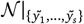.From this, it follows that we replace *v* in 𝒯 ^*^ by the correct cycle.

Since the blobtree 𝒯 ^*^ of 𝒩 was part of the sequence (𝒯_1_, …, 𝒯_*n* − 3_), exactly one of the candidate networks (𝒩 _1_, …, 𝒩 _*n* 3_) will be our original network 𝒩. By Frohn et al. (2024) we know that tf-quarnets are enough to encode a triangle-free semi-directed level-1 network. Hence, this will be the only network with a (weighted) tf-quarnet consistency score of 1 and so Squirrel returns the correct network 𝒩.

Whereas in Theorem 1 we showed that Squirrel reconstructs a network from its set of tf-quarnets, in practice, the input will not be a set of tf-quarnets coming from one unique network. In the following lemma we prove that even if the input tf-quarnets do not come from one unique network, Squirrel is still guaranteed to return a triangle-free semi-directed level-1 network.

### Lemma A.6.

*Let* 𝒬 *be a dense set of weighted tf-quarnets on* χ, *then* Squirrel *applied to* 𝒬 *returns a triangle-free semi-directed level-1 network on* χ.

*Proof*. Step A (Algorithm 1) of Squirrel relies on algorithms from Berry and Gascuel (2000) and Grünewald et al. (2009). It follows from those papers that Algorithm 1 always returns a sequence of phylogenetic trees on χ. To prove the lemma, we now need to show that Step B of Squirrel (Algorithm 2) returns a triangle-free semi-directed level-1 network on χ for any phylogenetic tree 𝒯 on χ. Since Algorithm 2 only alters the tree 𝒯 by replacing internal vertices of degree at least 4 by cycles with reticulations, the resulting network will always be level-1 and triangle-free. It remains to prove that the resulting network is a *valid* semi-directed network (i.e. a network that has a valid root-location, or equivalently, a network with no two reticulations oriented towards each other or towards the optional outgroup).

Clearly, when the first high-degree node is replaced by a cycle in the first iteration of the algorithm, there always is a location for the reticulation vertex that results in a partial network with a valid root-location. We can thus inductively assume that at the start of the other iterations, when we want to replace some internal vertex *v* by a cycle, the partial network constructed in the previous iteration has a valid root-location at some edge (or in case of a specified outgroup, the specific edge incident to the outgroup). Hence, the partial network can be rooted at this edge, forming a non-binary rooted phylogenetic network. We can then expand the high-degree node in this directed acyclic graph that corresponds to *v* in such a way that the resulting rooted phylogenetic network remains a directed acyclic graph. Thus, when we replace *v* by a cycle in the partially constructed semi-directed network, there is a location for the reticulation such that the partial network remains valid, which proves the lemma.

## B Random network generator

In this section we provide a concise description of the algorithm that was used to generate the random semi-directed level-1 networks in Section 2.1. The algorithm takes as input a number of leaves *n* and a number of reticulations *r* ≥ 1. In the special case that *r* = 0, we simply generate a random network with *r* = 1 reticulation and randomly turn it into a phylogenetic tree by deleting one of the reticulation edges (and suppressing the resulting degree-2 vertex). We first outline the general structure of the algorithm, while in the last paragraph we explain how the reticulation number is enforced. We emphasize that this algorithm is able to generate any semi-directed level-1 network on *n* leaves and with *k* reticulations, up to the labeling of the leaves.

The algorithm starts by building a random (possibly non-binary) phylogenetic tree on *n* leaves. In particular, it first generates a random tree on *n* vertices (by constructing a random spanning tree on a complete graph with *n* vertices), after which it attaches one leaf to every vertex. By suppressing all degree-2 vertices this results in a (possibly non-binary) tree with *n* leaves. Such a tree is transformed into a triangle-free semi-directed level-1 network by replacing every internal vertex of degree at least 4 by a random cycle. Finally, a random vertex of the cycle is assigned as a reticulation (while ensuring the resulting semi-directed network has a valid root location).

To enforce the correct reticulation number *r* in the resulting network, we ensure that the initial tree has exactly *r* internal vertices of high degree (i.e. a degree of at least 4). To this end, we iteratively adjust the tree until it satisfies this condition. In particular, if our initial tree has more than *r* high degree internal vertices, we contract the unique path between two random high degree internal vertices (without another high degree vertex on this path). This decreases the number of internal vertices with high degree by exactly one. On the other hand, if our initial tree has less than *r* of these high-degree internal vertices, we contract a random edge between two degree-3 nodes to create a new high degree vertex. This process is repeated until the tree has exactly *r* high-degree internal vertices.

